# One-pot synthesis of prenylated proteins utilizing *E. coli* cell-free expression

**DOI:** 10.1101/2022.03.03.482798

**Authors:** Lei Kai, Sonal, Tamara Heermann, Petra Schwille

## Abstract

Bottom-up synthetic biology is a powerful tool for uncovering the mechanisms underlying vital biological processes, such as signaling and cell polarization. The core principle of reconstituting cellular functions in their minimal forms can be achieved through modular protein design. However, assembling multiple purified proteins into a functional and synchronized system remains a technical challenge. The fact that many regulatory proteins show direct or indirect membrane interactions further exacerbates the complications. Here, we introduce the Cell-Free prenylated Protein Synthesis (CFpPS) system which enables the production of prenylated proteins in a single reaction mix, through reconstituted prenylation machinery. Not only does the CFpPS system offer a fast and reliable method for producing solubilized prenylated proteins, but it can also produce the protein of interest directly in the vicinity of biomimetic membranes, thus enabling microscopy-based functional assessment. As proof of principle, we demonstrate synthesis and solubilization of various important signaling proteins from the Ras superfamily, as well as membrane binding and extraction of the key polarity regulator Cdc42. Furthermore, our method can be used to confer membrane affinity to any protein, simply by adding a 4-peptide motif to the C-terminus of the protein. In sum, the CFpPS system offers a versatile and effective platform for designing peripheral membrane proteins for synthetic biology applications.

## Introduction

In their pursuit to understand the underlying principles of biological systems, scientists are usually challenged by the inherent redundancies in collective cellular functions. Hence, in recent years, attempts have been made to reduce cellular complexity and eventually deduce minimal functional requirements of life. This field, known as bottom-up synthetic biology, ultimately aims at the reconstitution of the simplest artificial system that mimics life ^1^. To this end, both biological and biology-inspired synthetic building blocks are enclosed in cell-sized compartments, consisting of self-assembled amphiphilic molecules, to create biomimetic or protocell-like systems ^2,3,4^. Successful reconstitutions of RNA synthesis and replication^1^, protein expression^5,6,7^, and *de novo* lipid synthesis machineries^8^ have been achieved using this approach. Boosted by advancements in biochemical and biophysical methodology, minimal cell research in recent years has been processing towards the reconstituting increasingly complex multi-protein and multimodule systems^9, 10^.

One technology that has greatly facilitated the assembly of multimodule systems is Cell-Free Protein Synthesis (CFPS)^6, 11^. CFPS systems host both the transcription and translation machineries, thus mimicking the cytoplasmic environment for protein production and facilitating a scale-up of functional modules in reconstituted systems. Recent advances in CFPS, in particular the well-established *E. coli*-based CFPS system, have led to a highly efficient and versatile platform for bottom-up synthetic biology^9, 11^. With the introduction of CFPS, a number of key features of cellular systems have been accomplished, including energy regeneration^12^, gene replication^13^, communication^14^, growth^15^, as well as the production of difficult proteins with post-translational modifications^16^. These merits of CFPS systems make them highly attractive tools for the construction of synthetic or minimal cell systems, which ultimately can self-replicate^13, 17^ or mimic cell division^7, 18^.

While CFPS systems are usually employed in a one-pot configuration, cellular processes often involve and require the presence of interfaces, such as the plasma membrane. Cell membranes do not only enable compartmentalization, but also localized homeostatic responses. For instance, small GTPases of the Rho and Ras families are important signaling nodes. These proteins display switch-like membrane binding allowing them to locally activate intracellular molecular pathways in response to diverse extracellular stimuli^19^. Small GTPases regulate various vital processes such as cell proliferation, differentiation, and apoptosis^19^. Malfunction of these proteins often results in serious or even life-threatening diseases. For example, K-Ras, H-Ras and N-Ras are oncogenic proteins, and missense mutations in Ras genes were found in nearly 25% of human cancers^20^. As an additional example, Cdc42 (a member of the Rho family) is an essential protein in eukaryotes, crucial for establishing cell polarity, intracellular trafficking, and regulation of cell growth^21^. Polarity establishment by Cdc42 relies on a positive feedback mechanism which is greatly facilitated by the switch-like membrane binding of Cdc42 ^22^. In sum, the highly diverse signaling pathways of Ras and Rho proteins heavily rely on their switchable membrane binding (Fig. 1a).

**Figure 1.**
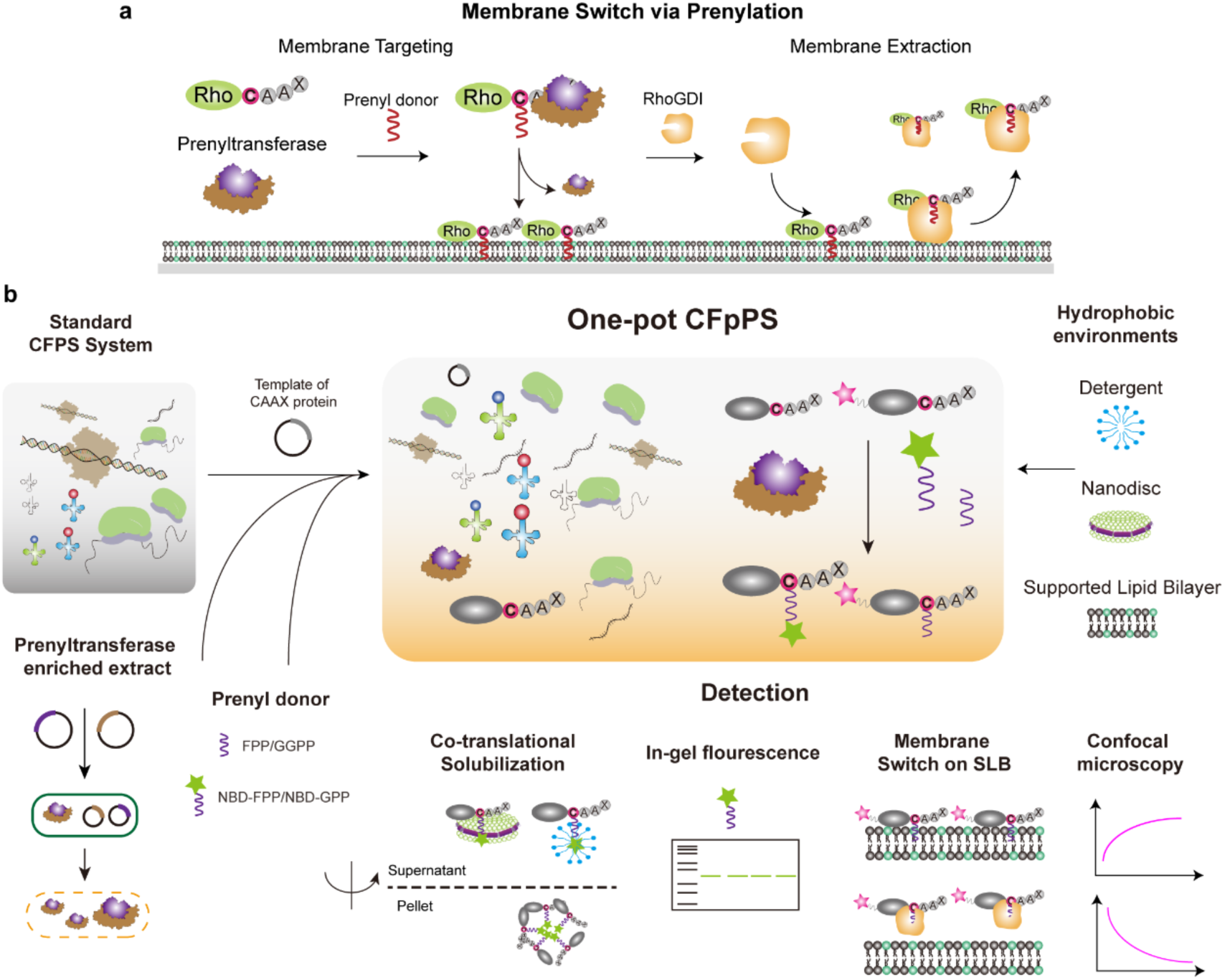
Reconstitution of switch-like membrane binding interactions using the cell-free prenylated protein synthesis (CFpPS) system. a Schematic illustration of reversible membrane binding of Rho proteins. Prenylation targets Rho proteins to the membrane in a switch-like manner. The Rho GDP dissociation inhibitor (RhoGDI) extracts GDP-bound prenylated Rho proteins from the membrane. b Co- translational prenylation was achieved by introducing prenyltransferase-enriched extract, the isoprenoid prenyl donor and the plasmid carrying the template of a target CAAX protein to the normal CFPS system. Newly expressed and prenylated proteins can either be directly incorporated into biomimetic membranes such as supported lipid bilayers (SLBs), or solubilized with amphipathic reagents, such as detergents or nanodiscs. Prenylation efficiency can be monitored in real-time by detecting the membrane recruitment of fluorescent fusion proteins on SLBs via confocal microscopy, or measured via in-gel fluorescence using fluorescent prenyl donors. Furthermore, for Rho proteins, RhoGDI can be introduced to extract proteins from the membranes.

The fast and dynamic shutting of small GTPases between the membrane and the cytosol is facilitated by a post-translational modification known as prenylation^23^. This enzymatic modification only occurs in eukaryotic cells and involves the irreversible addition of a lipid moiety to specific cysteine residues at the C-terminus of a protein^24^. The target proteins of prenylation are known as CAAX proteins for the conserved CAAX sequence motif at their C-termini, wherein “C” is the Cys that is being prenylated, “A” is an aliphatic amino acid and “X” can be any amino acid. Prenylation is catalyzed by enzymes known as prenyltransferases^25^. Depending on whether the isoprenoid involved is a 15-carbon farnesyl group or a 20-carbon geranylgeranyl group, prenyltransferases are known as farnesyltransferase (FTase) or geranylgeranyltransferase (GGTase), respectively; the corresponding lipid substrates are farnesyl-diphosphate (FPP) and geranylgeranyl- diphosphate (GGPP), respectively. Overall, prenyltransferases consist of four members: FTase^26^, and GGTase type-I^27^, type-II and type-III^28^. Mammalian FTase and GGTase-I are heterodimers, with an alpha subunit in common, and a unique beta subunit, which recognizes different CAAX sequences^29^. GGTase-II, also known as Rab GGTase^30^, requires an additional escort protein to mediate the binding to its target protein, and is mechanistically different from the other transferases, transferring two geranylgeranyl groups to the two cysteine residues at the C-termini of Rab proteins^31^. GGTase-III was recently reported with only one confirmed protein substrate Ykt6^28^. In contrast, FTase and GGTase-I have been well-studied with regard to their structures, functions and physiological roles. As a proof of concept, this study focuses only on the reconstitution of the activities of FTase and GGTase-I.

While dynamic membrane interactions are functionally important, hydrophobic domains may also compromise protein stability in hydrophilic environments, thus hindering reconstitution studies. The challenges of protein purification could potentially be bypassed by resorting to CFPS systems for prenylated proteins. Although prenylated CAAX proteins have previously been produced using the insect cell (sf21) CFPS system, the expression yield remained relatively low; it is likely that the availability of the prenylation machinery was limited by the low efficiency of preserving microsome fractions in the extract^32^. On the other hand, the prokaryotic *E. coli*-based CFPS system is considerably more efficient and cost-effective than eukaryotic CFPS systems for recombinant protein production; however, prokaryotes lack a prenylation machinery. Recent advances on the prenylation mechanism^33^, prenyl donor^34, 35^ and detection methods^36, 37^ have contributed to synthesize prenylated proteins directly within the *E. coli* CFPS system with both high yield and modification efficiency. Note that developing a reliable system for the production of prenylated proteins may be of even broader interest to synthetic biologists as switch-like membrane interactions hold immense potential for the modulation of simplified biological networks^38, 39^. Any protein could potentially be targeted to membranes by introducing a CAAX motif and employing the prenylation machinery.

Here, we establish a <u>c</u>ell-free <u>prenylated p</u>rotein <u>s</u>ynthesis (CFpPS) system by integrating the eukaryotic prenylation machinery into the *E. coli-*based CFPS system. We have produced extracts enriched with prenylation enzymes, systematically investigated the functionality of these extracts, and successfully demonstrated the co-translational prenylation of a range of representative proteins—Kras, Hras, RhoA, RhoC, Rac1 and Cdc42 from the Rho family—thus establishing a cell-free system for synthesis of prenylated proteins. There are two major accomplishments of this streamlined and integrated system: (i) the functionalization of the *E. coli*-based CFPS system, via enrichment with prenyltransferases, to express and prenylate CAAX proteins (ii) the co-translational solubilization of prenylated protein by introduction of different solubilizing agents such as detergents or lipid-based scaffolds. As proof of principle, we use the CFpPS system to reconstitute the membrane targeting and extraction of the eukaryotic polarity hub Cdc42 in a supported lipid bilayer (SLB), thus demonstrating the potential of such systems for bottom-up assembly of key biological processes such as cell polarization.

## Results

### Establishing the cell-free prenylated protein synthesis (CFpPS) system

For adding the prenylation machinery to the bacterial CFPS system, we prepared extracts from *E. coli* cells overexpressing a prenyltrasferase of interest (Supplementary Fig. 1b). The resulting combinations of the standard bacterial CFPS system with prenyltransferase- enriched extracts will, henceforth, be referred to as the CFpPS system (Fig. 1b). We selected FTase and GGTase-I from rat since these prenyltransferases express well in *E. coli* and are straightforward to purify (Supplementary Fig. 1-3)—features that have previously facilitated their *in-vitro* characterization ^40^. To validate the efficiency of the CFpPS system, we carried out prenylation with a fluorescently labeled lipid substrate (Fig. 2b) and detected the prenylated protein on SDS-PAGE gels using in-gel fluorescence (Fig. 1b). To serve as protein substrates for prenylation, we designed different chimeric proteins: The C-terminal amino acids of the small GTPases KrasB and Cdc42 were added to the C-terminus of the well-characterized Glutathione S-Transferase (GST) protein, connected by a rigid helical linker (Fig. 2a and Supplementary methods). FTase catalyzed the prenylation of GST-CAAX_KrasB_ by NBD-GPP, whereas GGTase-I catalyzed prenylation of GST- CAAX_Cdc42_ with NBD-FPP. The combination of these model reaction components enabled us to optimize parameters for the CFpPS system, such as concentrations of isoprenoids and the ratio of the component extracts for subsequent experiments.

**Figure 2.**
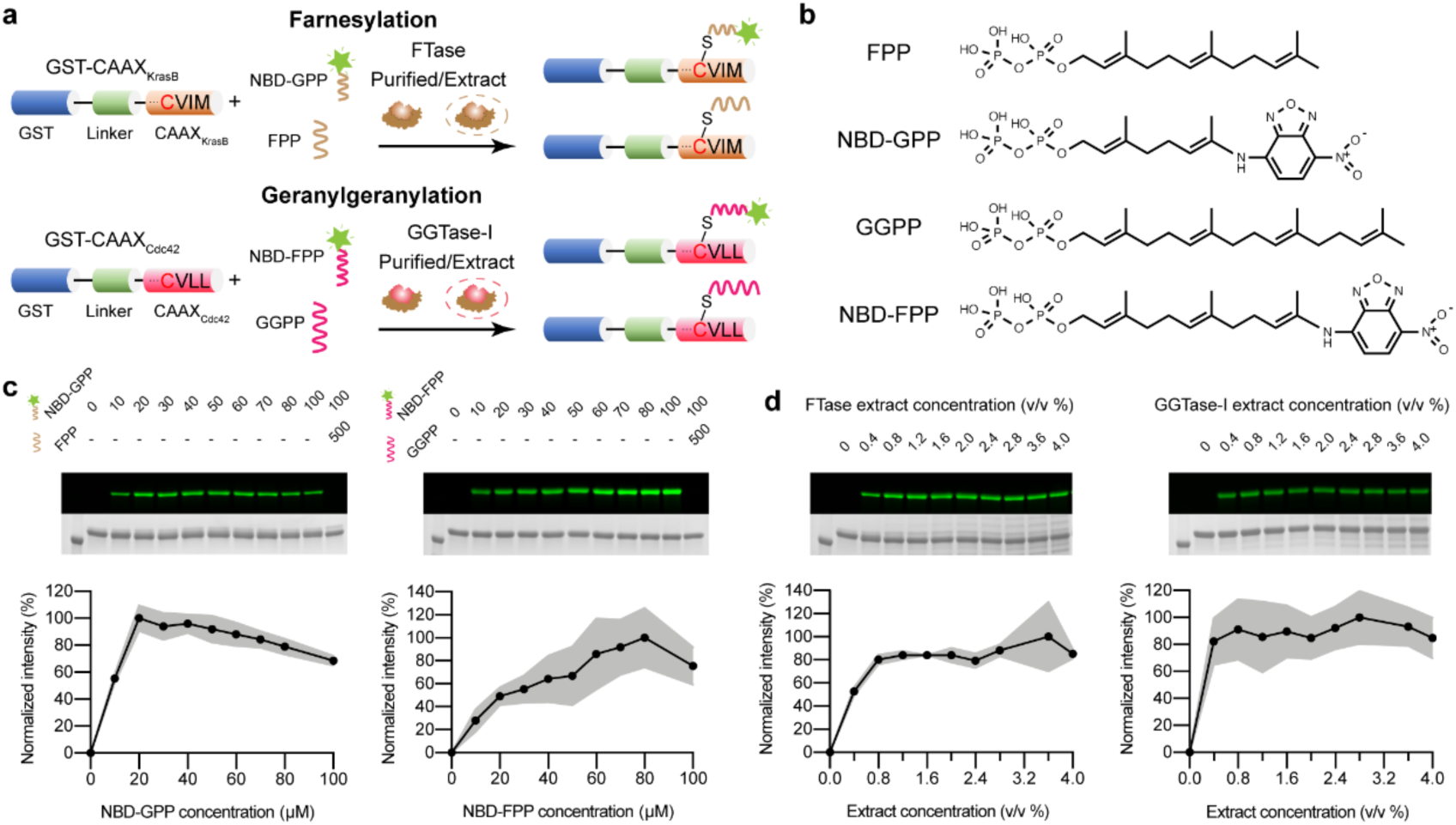
Concentration optimization of fluorescent prenyl donor and prenyltransferase-enriched extracts for *in vitro* prenylation reactions. a Schematic illustration of the chimeric proteins GST- CAAX_KrasB_ and GST-CAAX_Cdc42_ that are prenylated via FTase or GGTase-I, respectively. b Chemical structures of lipid donors. c Titration of the NBD-modified prenyl donors with purified prenyltransferases using in-gel fluorescence. NBD-GPP and FTase were used for the farnesylation of 10 µM GST-CAAX_KrasB_ (Left panel); NBD-FPP and GGTase-I were used for the geranylgeranylation of 10 µM GST-CAAX_Cdc42_ (right panel). Last lane of each gel shows the competition assay performed by adding the unlabeled analogue—either FPP or GGPP—at a concentration 5-fold of the highest tested for each NBD-modified analogue. Concentration (µM) of prenyl donors in each reaction is stated above the corresponding gel lane. d Titration of prenyltransferase-enriched extracts using In-gel fluorescence: FTase-enriched extract was used for the farnesylation of 10 µM GST-CAAX_KrasB_ with 20 µM NBD-GPP (left panel); GGTase-I-enriched extract was used for the geranygeranylation of 10 µM GST-CAAX_Cdc42_ with 80 µM NBD-FPP (Right panel). Extract concentration is shown as percentage volume of prenyltransferase-enriched extract included in the standard *E. coli* CFPS. Each image panel includes a representative gel imaged in fluorescence mode to visualize NBD (upper one) and colorimetric mode to visualize Coomassie staining (lower one). In all graphs, mean values from three independent replicates are shown as black dots, while the grey shading represents standard deviation, n=3.

**Figure 3.**
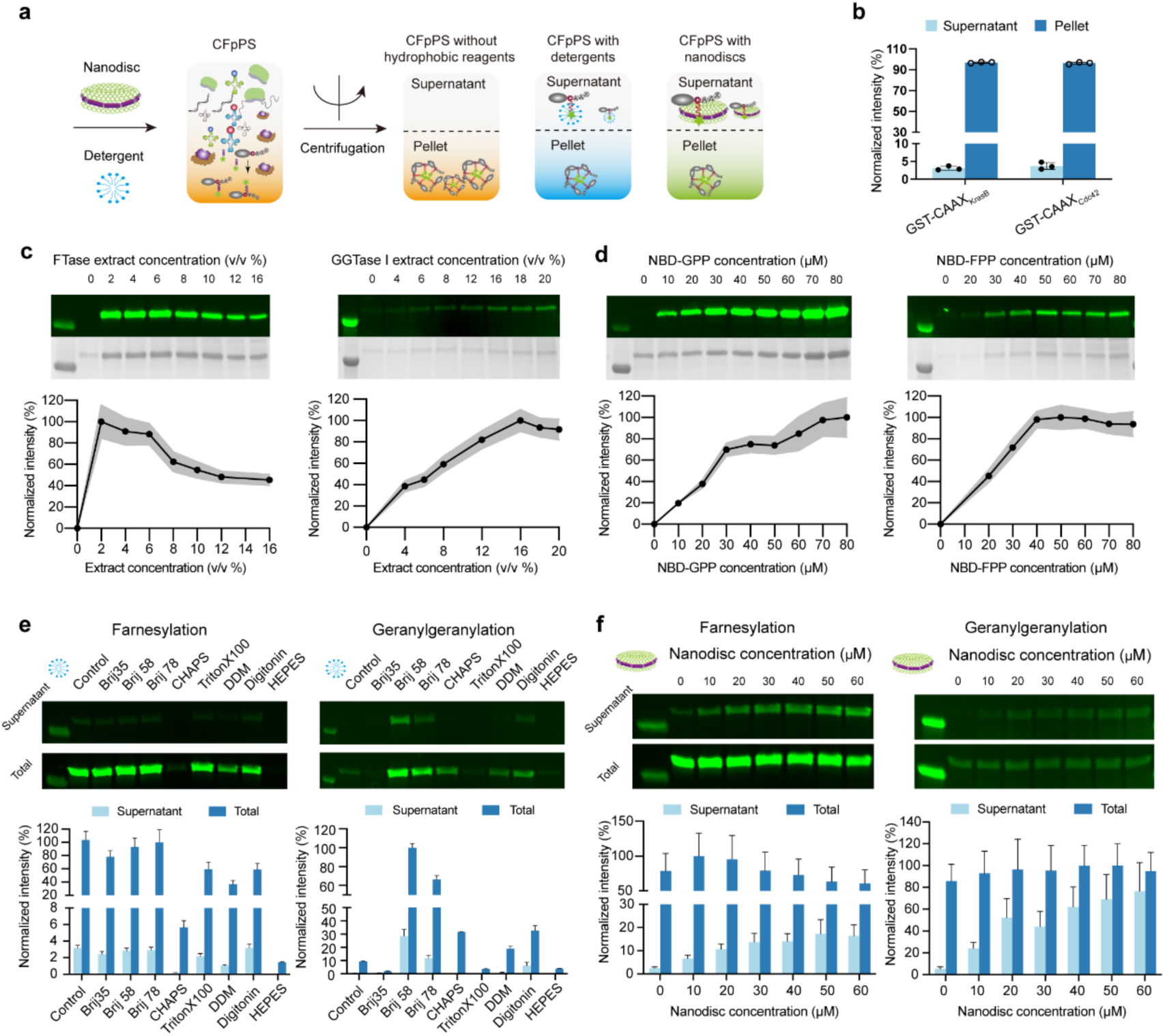
Co-translational expression and solubilization of prenylated protein by CFpPS. **a** Schematic depicting the expression and solubilization of prenylated CAAX proteins in CFpPS systems with or without solubilizing additives. **b** Prenylated GST-CAAX_KrasB_ or GST-CAAX_Cdc42_ demonstrate low solubility after co-translational prenylation in CFpPS extracts lacking solubilizing additives. 20/80 µM NBD- GPP/FPP were used and prenylated proteins were measured using in-gel fluorescence in the supernatant and the pellet fractions after centrifugation at 20,000xg. Measurements were normalized to the mean total protein amount in both the pellet and soluble fractions for each protein. Symbols represent intensity measured in 3 independent replicates. **c** Concentration optimization of prenyltransferase-enriched extract in CFpPS system using in-gel fluorescence: (Left) FTase-enriched extract; (Right) GGTase-I-enriched extract. Extract concentration is shown as percentage volume of prenyltransferase-enriched extract included in the standard *E. coli* CFPS. **d** In-gel fluorescence analysis for optimizing the concentration of NBD-modified prenyl donor in the CFpPS system. Panels **c** and **d** include representative gels imaged in fluorescence mode to visualize NBD (upper ones) and colorimetric mode to visualize Coomassie staining (lower ones). Pellet fractions from CFpPS reactions were used in the gels. Mean values of three independent replicates are shown as black dots, while the grey shading represents standard deviation. **e** Screening of detergents for soluble expression of farnesylated GST-CAAX_KrasB_ (left) and geranylgeranylated GST-CAAX_Cdc42_ (right). Respective control reactions were performed without any detergent. All intensity values were normalized using the highest averaged values. **f** Nanodisc titration for the soluble expression of GST-CAAX_KrasB_ (left) and GST-CAAX_Cdc42_ (right) in CFpPS system. Fluorescence intensities of the protein band for each fraction were measured through in-gel fluorescence and normalized to the highest averaged value.

Before testing prenylation extracts, we examined whether the purified prenyltransferases FTase and GGTase-I could prenylate their purified protein substrates GST-CAAX_KrasB_ and GST-CAAX_Cdc42_, respectively (Fig. 2c; Supplementary Fig. 2-3 for the purification of prenyltransferases). This enabled us to validate the in-gel fluorescence assay and to determine the effective concentrations of NBD-modified prenyl donors in a more controlled setting. For both reactions, at a fixed concentration of 10 µM protein substrate, the fluorescence intensity of the protein substrate’s band increased with ascending concentrations of its isoprenoid substrate, up to a certain concentration. Maximum activity was achieved at 20 µM isoprenoid concentration for farnesylation and 80 µM for geranylgeranylation. These were the concentrations chosen for subsequent tests with extracts. In both cases, prenylation output decreased marginally when isoprenoid concentration was further increased, possibly due to concentration-dependent aggregation of the isoprenoid substrates. Lastly, the protein band showed no fluorescence when natural isoprenoid substrates FPP and GGPP were included in the reactions at five times the highest concentration of their fluorescent analogues NBD-GPP and NBD-FPP, respectively (Fig. 2c, right-most lanes in gels). This competition assay confirmed that read- outs of in-gel fluorescence were specific to the prenylation reactions of interest.

Next, we used in-gel fluorescence to verify that prenylation activity was intact in prenyltransferase-enriched cell extracts. For enriched extracts, competent *E. coli* cells were simultaneously transformed with plasmid vectors carrying the alpha and beta subunit of the prenyltransferase of interest (either FTase or GGTase-I; Fig. 1), and overexpression was induced using IPTG. After induction, crude extract was prepared using the standard S30 preparation procedure. Using previously optimized concentrations of the fluorescent isoprenoids, the functionality of each prenylation extract was verified through in-gel fluorescence read-outs (Fig. 2d). Prenylation outputs saturated at around 0.8% (v/v) for both farnesylation extract and geranylgeranylation extracts, which roughly corresponds to 0.83 µM purified FTase and 1.22 µM GGTase-I (Supplementary Fig. 4).

Finally, to complete the set of components desired in the CFpPS system, instead of using purified protein substrates, we allowed the chimeric substrates to be directly expressed in the prenylation extracts. As expected, the prenylated form of either GST-CAAX_KrasB_ or GST-CAAX_Cdc42_ ended up in the pellet fraction upon centrifugation due to the increase in their hydrophobicity (Fig. 3b). We thus resolved the pellet fraction on SDS-PAGE for subsequent in-gel fluorescence assays to titrate the composition of the CFpPS system. First, we optimized the ratio of prenyltransferase-enriched extract to standard CFPS extract to achieve a maximum prenylation output. This screen was carried out at a fixed NBD-modified isoprenoid concentration of 30 µM (Fig. 3c-d). Second, we used the extract ratios of maximum activity (2% v/v for FTase extract and 16% v/v for GGTase-I extract) to titrate the NBD isoprenoid concentrations. Unlike the reaction with defined components, the reaction in the CFpPS system saturated at higher concentrations of NBD-lipid donors (e.g., 80 µM instead of 20 µM NBD-GPP for farnesylation). In addition, more enriched extract was required to achieve the same prenylation output. The reduction in the CFpPS system’s efficiency observed when protein substrates were co-translationally prenylated could be due to multiple reasons. For instance, non-specific interactions with residual membrane vesicles from the extract^41, 42^ could limit the availability of either the lipid donors or the prenyltransferases, and hence lead to higher saturation concentrations of both substrate and enzymes.

### Soluble expression of prenylated proteins

As shown in Figure 3b, the solubility of prenylated CAAX proteins was greatly decreased due to the hydrophobicity of the prenyl group. Having obtained basic parameters for co-translational prenylation, we sought to improve the soluble expression of prenlyated proteins for downstream functional analysis by introducing solubilizing reagents to the CFpPS system (Fig. 3a). First, we tested detergents, which are often used in CFPS systems to improve the solubility of expressed membrane proteins^43^. The compatibility of seven commonly used detergents were evaluated by determining prenylation efficiency in the presence of the detergent using in-gel fluorescence (Supplementary Fig. 5). Next, these detergents were directly introduced in the CFpPS system to assess their ability to keep the prenylated protein soluble. After centrifugation, supernatant, pellet, and total protein fractions were collected and evaluated for the presence of the prenylated proteins (Fig. 3e). Nearly all detergents tested showed little increase in the solubility of farnesylated GST-CAAX_KrasB_ (Fig. 3e). In contrast, the solubility of the geranylgeranylated GST- CAAX_Cdc42_ increased considerably in the presence of certain detergents such as Brij58 (Fig. 3e right panel). Interestingly, the introduction of detergents could also improve the total prenylation efficiency for GGTase-I, with a nearly 10-fold increase in the presence of Brij58.

As an alternate solubilization strategy, particularly for farnesylation, we introduced nanodiscs to the CFpPS system. Nanodiscs are discoidal lipid bilayers stabilized by the presence of amphipathic protein belts and are thus closer mimics of the cell membrane^44^ (See Supplementary methods for details). Negatively charged lipids were included in the nanodiscs to promote the association of prenylated proteins through the conserved polybasic regions of native CAAX proteins^45^. Nanodiscs could in fact increase the soluble fraction of farnesylated GST-CAAX_KrasB_ up to 6-fold, although there was a slight decrease in total modified proteins (Fig. 3f). For geranylgeranylated GST-CAAX_Cdc42_, nanodiscs could increase the soluble fraction approximately 14-times compared to the control, corresponding to 72% of the total modified protein. Notably, the total modified protein stayed relatively stable for geranylgeranylation (Fig. 3f right panel). In summary, nanodiscs could greatly improve the solubility of farnesylated GST-CAAX_KrasB_, while either the detergent Brij 58 or nanodiscs promoted the solubility of geranylgeranylated GST- CAAX_Cdc42_.

Finally, we tested the performance of the optimized CFpPS system in expression and solubilization of a broader representative CAAX proteins. The CFpPS system succeeded in expression and prenylation of not just chimeric proteins with CAAX sequences, but also a selection of the native small GTPases; furthermore, all proteins except RhoA and RhoC could be produced in a soluble form, aided either by nanodiscs or by Brij 58 (Table 1, Supplementary Fig. 6).

**Table 1.** Summary of the CFpPS system’s performance in expressing and solubilizing chimeric CAAX-proteins and native small GTPases.

| Constructs | C-terminal sequences | Type of | Detected | Soluble |
| --- | --- | --- | --- | --- |

|  |  | Prenylation | Modification | Modification |
| --- | --- | --- | --- | --- |
| GST-CAAX<sub>KrasB</sub> | GKKKKKKSKTKCVIM | F/G | + | + |
| GST-CVIM | GCVIM | F/G | + | + |
| GST-CAAX<sub>Cdc42</sub> | KKSRRCVLL | G | + | + |
| KrasB | GKKKKKKSKTKCVIM | F/G | + | + |
| Hras | CVLS | F | + | + |
| Cdc42 | KKSRRCVLL | G | + | + |
| RhoA | RRGKKKSGCLVL | G | + | - |
| RhoC | KNKRRRGCPIL | G | + | - |
| Rac1 | KKRKRKCLLL | G | + | + |

F, farnesylation; G, geranylgeranylation.

### Chimeric proteins synthesized by the CFpPS system bind to biomimetic membranes

In a cell, prenylation of small GTPases is often followed by additional post-translational modification steps, such as methylation of the CAAX cysteine and cleavage of the last three amino acids^24^. Enzymes for these steps are currently not included in the CFpPS system. We therefore wanted to test whether prenylated proteins produced through CFpPS show membrane interaction. To this end, we leveraged the fact that the CFpPS system that it can easily be carried out in the vicinity of biomimetic membranes and membrane binding can be monitored using confocal microscopy.

Co-translational prenylation of mCherry-CAAX_Cdc42_ was carried out by the CFpPS system on top of a planar supported lipid bilayer (SLB). A confocal microscopy-based assay was applied to monitor membrane targeting of mCherry-CAAX_Cdc42_ (Fig. 4a). Localization of the protein substrate could be followed using the fluorescence of mCherry, whereas a small fraction of the fluorescently labeled lipid Atto-488 PE was included in the SLB to visualize the membrane. The lipid composition of the SLB was identical to the one used previously for nanodiscs. All components of the CFpPS reaction, except the prenyl donor GGPP, were mixed in a chamber with a preformed SLB. GGPP was withheld to time the triggering of the prenylation process. As expected, upon addition of GGPP, we observed an increase in mCherry fluorescence on the SLB, indicating that the geranylgeranylated mCherry- CAAX_Cdc42_ was attaching to the membrane. Orthogonal sections (Fig. 4b) through the confocal stack as well as measured Z-axis intensity profiles (Fig. 4c) show that the mCherry fluorescence occupies the same plane as the membrane. The mCherry signal continued to increase until it saturated roughly 15 minutes after the addition of GGPP. Thus, including the CAAX motif in an otherwise soluble protein is sufficient to target it to a membrane using the CFpPS system.

**Figure 4.**
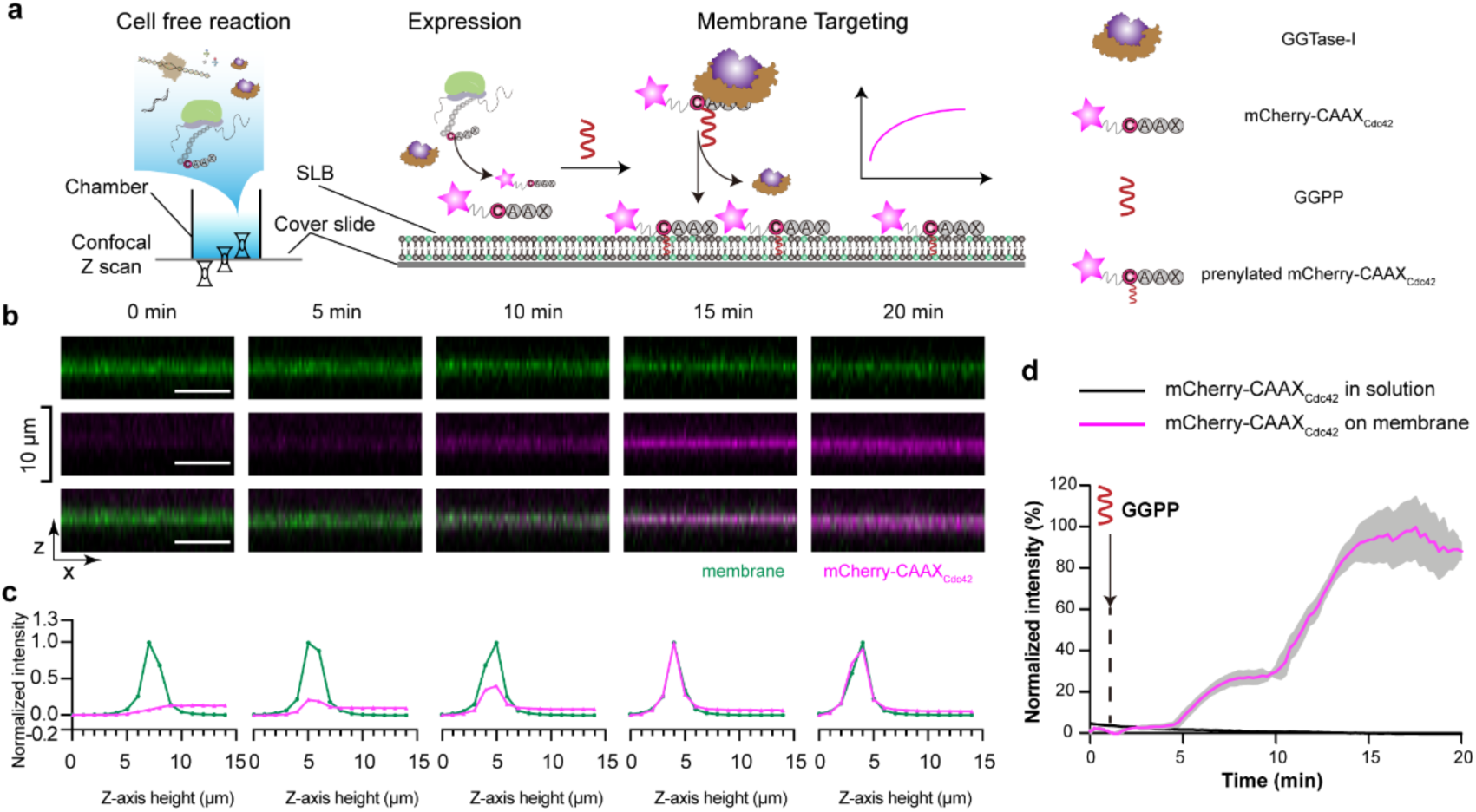
Prenylated mCherry-CAAX_Cdc42_ produced using the CFpPS system binds to biomimetic membranes. **a** Schematic illustration of the membrane targeting of mCherry-CAAX_Cdc42_, as studied on supported lipid bilayers (SLBs) using confocal microscopy **b** Orthogonal views of the SLB membrane (upper, green), mCherry-CAAX_Cdc42_ (middle, magenta) and a merge of both channels (lower) at different time points after prenylation was initiated in the CFpPS reaction by adding GGPP. The SLB composition is 80% DOPC, 19.95% DOPS, 0.05% Atto-488 PE. All scale bars are 10 µm. **c** Normalized intensities of corresponding images from panel **b**. Intensities of mCherry-Cdc42 were normalized to the maximum and minimum intensities recorded in z-stack during the time lapsed experiments; intensities of membrane channel were normalized to the maximum and minimum intensities recorded in z-stack in each time point. **d** Time series of mCherry-CAAX_Cdc42_ intensity on the membrane (pink) and in solution (black). Intensities were normalized to the maximum and minimum intensities measured during a time-lapse experiment. Solid lines represent the mean intensity measured over a 75-pixel by 75-pixel region and grey shading represents the standard deviation. The displayed data is representative for the 3 independent replicates.

### Cdc42 synthesized by the CFpPS system shows reversible membrane binding and native protein interactions

After validating the membrane binding of chimeric mCherry-CAAX_Cdc42_ upon prenylation, we tested whether the reversible membrane binding of the full-length Cdc42 could be reconstituted using the CFpPS system. Prenylated Cdc42 is known to bind to negatively charged lipids and this charge preference could indeed be confirmed using purified Cdc42 (Supplementary Fig. 7). Additionally, Cdc42 can be extracted from the membrane by the Rho GDP dissociation inhibitor (RhoGDI) ^46^, which sequesters the hydrophobic geranylgeranyl moieties of prenylated Cdc42. Following a similar one-pot experimental design as that used for the chimeric CAAX protein, an N-terminal mCherry fluorescent tag was fused to full-length Cdc42 (Fig. 5a) for confocal imaging. The membrane targeting process was initiated via the addition of GGPP, while the membrane extraction process was primed upon addition of RhoGDI (Fig. 5a, d, e). As with the mCherry-CAAX_Cdc42_, the mCherry-Cdc42 intensity on the membrane increased until saturation at roughly 15 minutes since the addition of GGPP (Fig. 5d). Orthogonal view (Fig. 5b) and Z-axis intensity profiles (Fig. 5c) both showed that the localization of mCherry-Cdc42 had a significant overlap with the membrane at that time-point.

**Figure 5.**
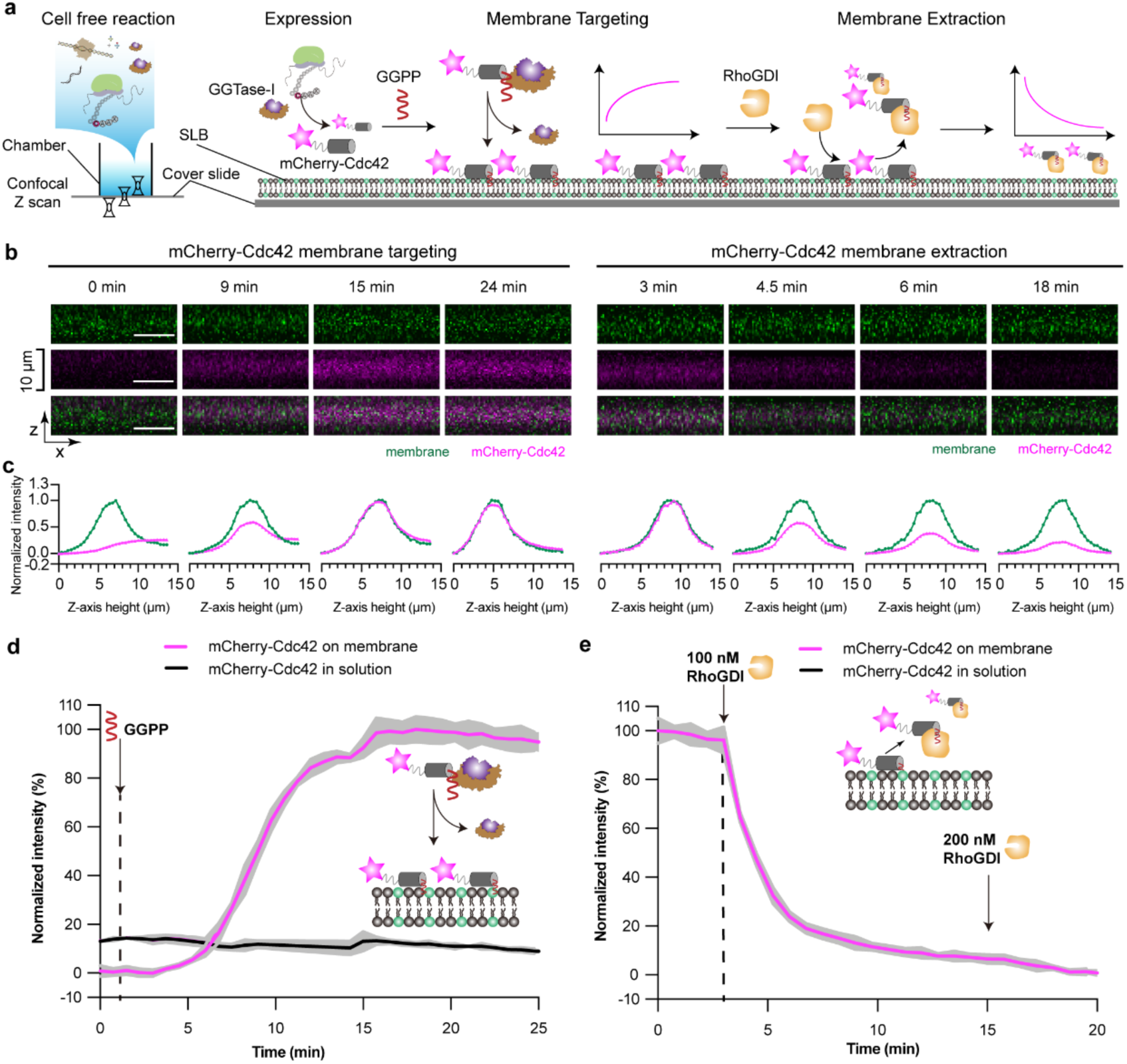
mCherry-Cdc42 produced using the CFpPS system shows expected membrane binding and RhoGDI-dependent extraction. **a** Schematic illustration of the reconstitution of Cdc42’s membrane targeting and RhoGDI-dependent membrane extraction on SLBs, as visualized using confocal microscopy. **b** Representative images of orthogonal views of the SLB membrane (upper, green), mCherry-Cdc42 (middle, magenta) and a merge of both channels (lower) at different time points. Time was measured from the addition of GGPP during membrane targeting and from the initiation of time-lapse imaging for the extractions process. All scale bars denote 10 µm. **c** Normalized intensities of corresponding images from panel **b**. Intensities of mCherry-Cdc42 were normalized to the maximum and minimum intensities recorded in z-stack during the time lapsed experiments; intensities of membrane channel were normalized to the maximum and minimum intensities recorded in z-stack in each time point. During extraction, the same normalization process was followed for the membrane channel; while, the intensities of mCherry- Cdc42 was normalized to the maximum intensity recorded in the z-stack at 3 minutes before the addition of RhoGDI. **d-e** Time series of mCherry-Cdc42 intensity on the membrane (pink) and in solution (black) during membrane targeting (**d**) and membrane extraction (**e**). Intensities were normalized to the maximum and minimum intensities measured during a time-lapse experiment. Solid lines represent the mean intensity measured over a 75-pixel by 75-pixel region and grey shading represents the standard deviation. The displayed data is representative for the 3 independent replicates. The lipid composition of SLBs used in this figure is 80% DOPC, 19.95% DOPS, 0.05% Atto-488 PE.

To verify the specificity of mCherry-Cdc42’s membrane binding, we then added RhoGDI to trigger the membrane extraction of Cdc42^47^. Upon addition of RhoGDI, mCherry fluorescence sharply decreased on the membrane, indicating that membrane-bound mCherry-Cdc42 was extracted from the SLB (Fig. 5 b-c, Fig. 4e). After nearly two-thirds of the membrane-bound Cdc42 had been extracted, introducing additional RhoGDI did not lead to further decrease in mCherry fluorescence on the membrane indicating that the remaining Cdc42 may not be extractable (Fig. 5e). Unlike the full-length Cdc42, the chimeric construct mCherry-CAAX_Cdc42_ could not be extracted by RhoGDI after binding to the SLB, confirming that the extraction process we saw for the full-length Cdc42 results from specific protein-protein interactions (Supplementary Fig. 8). Altogether the RhoGDI experiments demonstrate that proteins synthesized by the CFpPS system not show expected membrane binding, but can also retain their native interactions with other proteins. Furthermore, mCherry-Cdc42 purified after production in the CFpPS system showed a similar membrane targeting behavior and specific extraction by RhoGDI, indicating that none of the components of CFpPS participate in the membrane binding (Supplementary Fig. 9). In summary, the CFpPS system constitutes a suitable platform for direct reconstitution of dynamic membrane binding of Cdc42 on a biomimetic membrane environment provided by the SLB.

## Discussion

In this study, we have successfully established the combined expression and membrane targeting of proteins by leveraging CAAX prenylation in an *in vitro* cell-free setup. We have achieved this by combining the standard cell-free protein synthesis (CFPS) system with prenylation machineries corresponding to different lipid substrates. The resulting prenyltransferase-enriched cell-free protein system, referred to as CFpPS, enabled the efficient expression and co-translational prenylation and solubilization of both synthetic constructs with exogenous CAAX motifs and natural CAAX proteins. Furthermore, using the polarity hub Cdc42 as a test case, we could demonstrate that the prenylated CAAX protein produced from CFpPS extracts shows expected interactions with lipid membranes, as well as other proteins (RhoGDI), and can directly be reconstituted in biomimetic membranes for subsequent analysis.

Many important signaling proteins exhibit binding to scaffolds such as cell membranes and thus often possess hydrophobic motifs. On the other hand, such motifs complicate the expression, purification and reconstitution of those proteins in their soluble form in hydrophilic environments as preconditions for bottom-up synthetic biology ^9^. In order to stabilize prenylated proteins in solution, previous *in vitro* studies probing regulatory interactions of CAAX proteins have often resorted to using either truncated versions without the C-terminus^48, 49^ or unmodified proteins^50, 51^. Other compensatory methods have included replacement of the C-terminus with hydrophobic helices^52, 53^, genomic engineering of host cells to co-express corresponding prenyltransferases^54^, and *in vitro* prenylation with purified prenylatransferase^34, 55^. However, removal or replacement of the C-terminus can create artifacts by perturbing the protein’s native membrane interaction^56, 57^, while the other approaches have disadvantages such as requiring tedious genomic engineering or resulting in low protein yields^54^. Suzuki *et al*. leveraged the endogenous prenyltransferases in an insect cell lysate to synthesize prenylated CAAX proteins^32^. Still, this method also suffers from low protein yields since, similar to *in vivo* approaches, the prenylation capacity is limited by the availability of endogenous prenyltransferase in the lysate’s microsome fraction. In addition, co-existence of both endogenous FTase and GGTase-I could potentially result in a heterogenous pool of proteins, since some Ras and Rho GTPases such as KrasB^58^ or RhoA^59^ can undergo both farnesylation and geranylgeranylation.

By combining the *E. coli*-derived CFPS system with prenyltransferase-enriched lysates, the CFpPS system successfully addresses all these challenges. First, the system is entirely cell-free and requires neither the modification of host organisms nor protein purification to perform *in vitro* prenylation reactions. Second, the use of bacterial lysates allows abundant supply of prenyltransferases and thus leads to considerably improved yields. After optimization, the CFpPS system could produce prenylated protein at a concentration of approximately 0.5 mg/ml. Cell-free reactions can also easily be scaled to meet yield requirements for different analyses^60^. Furthermore, the modular design of the system allows us to design prenylation reactions and substrates to suit our needs. One can choose extracts enriched with either farnesyl- or geranylgeranyl-transferase, or even their combination.

One major advantage of the CFpPS system is that solubilizing agents such as detergents or lipids can be introduced to co-translationally solubilize the prenylated CAAX-proteins. Absence of hydrophobic environments can lead to aggregation of the modified protein or unspecific binding to chromatography resins, leading to significant losses during protein purification^54^. Although detergents can reduce this unspecific binding, they can interfere with further functional characterization of the protein, e.g., by perturbing the integrity of biomimetic membranes when membrane binding or enzymatic activities are being tested^54^. For instance, when Tnimov *et al.* used a purified prenyltrasferase to modify RhoA and investigate its membrane delivery and extraction by RhoGDI, geranylgeranylated RhoA aggregated rapidly and was only soluble with stoichiometric amounts of GGTase-I^46^. In the CFpPS systems, we successfully solubilized both farnesylated and geranylgeranylated CAAX-proteins using either detergents or nanodiscs. Nanodiscs were highly effective in solubilizing farnesylated proteins, whereas the detergent Brij 58 was the most beneficial for soluble expression of geranylgeranylated proteins (Fig. 3e-f).

In the case of geranylgeranylation, we observed that certain detergents, such as Brij 58, Brij 78 and Digitonin not only improved solubility but also promoted an increase in the total amount of modified protein (Fig. 3e). More specifically, the presence of Brij 58 led to a 10- fold increase in total modified GST-Cdc42_CAAX_ and nearly 30% of the modified protein was accounted for in the soluble fraction. This effect was likewise detected in the *in vitro* geranygeranylation reaction; however, the improvement was small compared to that observed in the CFpPS system (Supplementary Fig. 5). Interestingly, the same effect was found for all selected Rho GTPases, such as Cdc42, RhoA, RhoC and Rac1 (Supplementary Fig. 6). Despite the overall improvement in the fraction of total modified protein, Brij 58 could only improve the solubility of Cdc42 amidst the prenylated Rho GTPases (Table 1 and Supplementary Fig. 6). It is possible that the improvement in modified protein fractions might be due to better solubility or availability of the prenyl donor GGPP in the presence of detergents. Our analysis suggests that solubilizing conditions would need to be optimized individually for each protein of interest. However, the CFpPS would also facilitate this process by offering an easy and low-cost platform for extensive screening of solubilizing agents.

Although detergents like Brij 58 could improve the solubility and yield of Rho GTPases such as Cdc42, downstream experiments with model membranes would still be impeded by the presence of detergents. To circumvent this impediment, we leveraged the fact that our CFpPS system integrates expression and geranylgeranylation in one compartment and included model membranes directly in this compartment. Successfully targeting the chimeric substrate mCherry-CAAX_Cdc42_ to a biomimetic membrane demonstrates that the CFpPS system can easily enable the membrane targeting of a soluble protein, simply by adding the 4-amino acid CAAX motif to the C-terminus of the protein sequence in a plasmid. Furthermore, targeting to the membrane could be timed by withholding the prenyl donor resulting in a switch-like membrane binding behavior, which can be highly beneficial for the study of signaling cascades. This property offers an added advantage over previously used reconstitution strategies such as including nickelated or biotinylated lipids in the membrane and adding a His tag or Strep tag, respectively, to the protein of interest^61, 62^.

A reversible membrane-binding switch is of great relevance to the functionality of the polarity regulator Cdc42^63, 64^. The CFpPS system successfully reconstituted the dynamic membrane interaction of Cdc42 on a supported lipid bilayer in a one-step reaction. Cdc42 produced by the CFpPS system not only shows the expected membrane binding, but could also be subsequently extracted from the membrane by RhoGDI (Fig. 5). Both membrane targeting and extraction processes were found to be similar to those observed for purified Cdc42 and GGTase-I (Fig. 5, Supplementary Fig. 9), indicating that proteins produced by the two methods were similar in quality and fully functional. The key advantage of the setup involving the CFpPS system atop an SLB is that additional protein regulators can easily be introduced into the same chamber. These proteins could be added in a purified form when their concentration is of key relevance. Alternatively, proteins of interest could be introduced as plasmids, which get translated alongside Cdc42 in the CFpPS system; this approach would consequentially open myriads of possibilities for easily testing chimeric proteins. Furthermore, considering the compatibility of microscopy analysis with the CFpPS setup, we offer a promising *in vitro* platform for the rigorous study of complex, multi- protein processes such as polarization.

In summary, we have successfully established the CFpPS system by including prenyltransferase-enriched cell extracts in the standard bacterial CFPS. The resulting CFpPS is capable of expression and prenylation of any protein containing a CAAX motif, and can produce solubilized membrane-binding proteins for various end uses and functional analyses. We postulate that the CFpPS method holds great potential for studying the elaborate protein interaction networks of small GTPases using bottom-up synthetic biology approaches.

## Material and Methods

### Bacterial strains and plasmids

A list of all plasmids generated and used in this study can be found in Supplementary Information (Supplementary Table 1). Protein and gene sequences as well as cloning procedures are provided in Supplementary Material and Methods.

### Protein expression and purification

Purifications of FTase and GGTase-I were performed according to previous protocols^34, 65^. Briefly, a single colony of *E. coli* Rosetta, carrying both plasmids for the *α* and the *β* subunit, was grown overnight at 30 ^°^C in 50 mL of Luria Bertani (LB) medium (10 g L^-1^ tryptone, 5 g L^-1^ yeast extract, 5 g L^-1^ NaCl), containing 50 μg/ml carbenicillin (CA), 50 μg/ml kanamycin (KAN) and 37 μg/ml chloramphenicol (CHL). Then, 500 mL terrific broth (TB) medium (20 g L^-1^ tryptone, 24 g L^-1^ yeast extract, 0.4% (v/v%) glycerol, 10% (v/v%) phosphate buffer (0.17 M KH_2_PO_4_/0.72 M K_2_HPO_4_)), including CA/KAN/CHL, was inoculated with 10 mL overnight culture and grown at 37 ^°^C until Abs_600_ was 0.9-1.0. Protein expression was induced by adding 0.4 mM IPTG and further incubation at 37 ^°^C for 4 hours (see Supplementary Fig. 1). Cells were harvested by centrifugation at 8,000xg at 4 ^°^C, followed by twice-washing steps with PBS buffer. The resulting cell pellet was resuspended in 30 mL lysis buffer (50 mM phosphate buffer pH 8.0, 0.3 M NaCl, and 1 mM TCEP, 0.1 mM phenylmethylsulfonylfluoride (PMSF)) and cells were disrupted via a single pass through a pre-cooled French Press at 17,000 psi prior to 30 min incubation on ice with 10 U/mL Benzonase nuclease. The lysate was then centrifuged at 20,000x g for 30 min. The supernatant was filtered through a 0.22 μm PVDF membrane and loaded onto the 5 mL HiTrap GST column via an ÄKTA pure protein system (Cytiva). Eluted peak fractions were pooled and 10-fold diluted with anion exchange buffer (50 mM phosphate buffer pH 8.0 and 1 mM TCEP). The resulting mixture was loaded onto an anion exchange Hitrap Q FF column (Cytiva) and washed with 50 column volumes of buffer (50 mM phosphate buffer pH 8.0 and 1 mM TCEP). FTase or GGTase-I were eluted through the linear increase of NaCl to a final concentration of 1 M. Peak fractions were collected and dialyzed against 5 L of dialysis buffer (50 mM HEPES, pH 7.2, 100 mM NaCl, 5 mM DTT). Dialyzed samples were further concentrated to 10 mg mL-1 using Amicon^®^ Ultra-15 Centrifugal Filter Unit (Merck Millipore). The final concentration of glycerol was adjusted to 50% (v/v%) for storage at -80 ^°^C.

GST-CAAX_KrasB_, GST-CAAX_Cdc42_ and RhoGDI were expressed in *E. coli* BL21 (DE3) in TB medium, following the same protocol as the FTase and GGTase-I purification, except that IPTG concentration was increased to 1 mM and protein expression was performed via overnight incubation at 16 ^°^C. GST-CAAX_KrasB_ and GST-CAAX_Cdc42_ were stored in storage buffer (50 mM HEPES, pH 7.8, 150 mM NaCl, 1 mM TECP, 15% v/v glycerol) and flash-frozen in liquid nitrogen prior to their storage at -80 ^°^C. For purification of RhoGDI via Ni-NTA column, the following buffers were used: lysis buffer (50 mM Tris-HCl, pH 7.4, 300 mM NaCl, 2 mM TCEP, 5 mM MgCl_2_, 0.1 mM PMSF), wash buffer (50 mM Tris-HCl pH 7.4, 300 mM NaCl, 2 mM TCEP, 5 mM MgCl_2_,10 mM imidazole) and elution buffer (50 mM Tris-HCl pH 7.4, 300 mM NaCl, 2 mM TCEP, 5 mM MgCl_2_, 250 mM imidazole). RhoGDI was further purified through gel filtration on a Superdex 75 size exclusion column (storage buffer: 50 mM Tris-HCl pH 7.4, 300 mM NaCl, 1 mM TCEP, 5 mM MgCl_2_) and protein fractions were pooled and concentrated prior to storage in single-use aliquots at -80 ^°^C.

mCherry-Cdc42 was expressed as outlined above. Purification was performed using HisTrap™ column and Superdex 75 column and the protein was stored in storage buffer (20 mM HEPES pH 7.8, 150 mM NaCl, 2 mM MgCl_2_, 0.1 mM GDP, 0.5 mM TCEP, 10% v/v glycerol) at -80 ^°^C as single-use aliquots.

### Preparation of S30 extract

Standard S30 extract for cell-free protein synthesis was prepared according to previously published protocols^66, 67^. Briefly, *E. coli* BL21 (DE3) cells were grown until mid-log growth phase (Abs_600_ around 3.0) in 2 L baffled Erlenmeyer flasks. Cells were fast-chilled for 10 min under ice cold water and harvested via centrifugation at 8,000xg for 15 min. Cell pellets were washed three times with pre-cooled S30 A buffer (10 mM Tris-acetate pH 8.2, 14 mM Mg(OAc)_2_, 60 mM KCl, 6 mM ß-mercaptoethanol) and resuspended with 110% (v/w%) volume S30 B buffer (10 mM Tris-acetate pH 8.2, 14 mM Mg(OAc)_2_, 60 mM KCl, 1 mM DTT and 1 mM PMSF). The resuspended cells were disrupted via a single pass through the French Press at 17,000 psi. The resulting lysates were clarified via two rounds of centrifugation at 30,000xg. The supernatant was mixed with 0.3 volume of pre-incubation buffer (300 mM Tris-acetate pH 7.6, 10 mM Mg(OAc)_2_, 10 mM ATP, 80 mM Phosphoenolpyruvat (PEP), 5 mM DTT, 40 μM each of the 20 amino acids, 8 U mL^-1^ pyruvate kinase) and incubated at 37 ^°^C for 80 mins. Samples were then dialyzed for 2 hours against the 100-fold volume of S30 C buffer (10 mM Tris-acetate pH 8.2, 14 mM Mg(OAc)_2_, 60 mM KOAc, 0.5 mM DTT) and again overnight at 4 ^°^C. The dialyzed samples were centrifuged at 30,000xg for 30 min, the supernatant was collected into small aliquots, frozen with liquid nitrogen and stored at -80 ^°^C until further usage. For the prenyltransferase-enriched extract, double transformed cells were induced at an Abs_600_ of 1.0 with 0.4 mM IPTG for 3 hours (final Abs_600_ was around 4.0). All further steps were identical to the outlined standard S30 extract preparation.

### *In vitro* prenylation assay and in-gel fluorescence analysis

For in vitro protein prenylation, reaction mixtures (volume 20 μL) were composed of: 10 μM CAAX protein, 0.4 μM FTase or 2 μM GGTase-I and the respective indicated concentrations of NBD-GPP or NBD-FPP in prenylation buffer (50 mM HEPES pH 8.0, 300 mM NaCl, 20 μM ZnSO_4_, 2 mM MgCl_2_, 0.5 mM TCEP, 100 μM GDP). Reaction mixtures were incubated for 2 hours at 25 ^°^C and quenched by adding 10 μL 4x Laemmli Sample buffer (Bio-Rad). Samples were boiled at 95 ^°^C for 5 min and each 8 μL were loaded onto 12% SDS-PAGE gels. Prenylated protein bands were visualized in gel using an Amersham Imager 600RGB (Cytiva) (excitation light: blue epi light (460 nm), emission filters: Cy2 (525BP20)). After fluorescent imaging, gels were stained with Instant Blue^TM^ (Expedeon) and scanned. For competition assays, natural prenyl-donors FPP or GGPP were additionally introduced beside the corresponding NBD analogues. Fluorescent images and Coomassie stained images were analyzed by Fiji^68^. Fluorescence intensities were calibrated by the densitometry from the respective Coomassie stained gel images to reduce loading error.

#### Cell-Free prenylated Protein Synthesis (CFpPS)

Cell free protein synthesis reactions were prepared according to protocols previously published by us and Kigawa *et al.*^66, 67^. In brief, a typical CFPS reaction contained: 55 mM Hepes-KOH buffer (pH 7.5), 1.7 mM DTT, 1.2 mM ATP (pH 7.0), 0.8 mM each of CTP (pH 7.0), GTP (pH 7.0), and UTP (pH 7.0), 80 mM Creatine phosphate (CP), 80 μg mL^-1^ Creatine kinase (CK), 2.0% (v/v%) PEG-8000, 0.65 mM 3,5-cyclic AMP, 68 μM folinic acid, 170 μg mL^-1^ E. coli total tRNA, 200-250 mM potassium glutamate, 27.5 mM ammonium acetate, 15-20 mM magnesium acetate, 2.0 mM of each of the 20 amino acids, 10 μg mL^-1^ T7 polymerase (prepared according to our previous protocol^67^, see Supplementary Methods), 30% (v/v %) S30 extract, 15 ng μL^-1^ plasmid template. A typical reaction volume was 50 μL and the reaction mixture was incubated at 30 ^°^C for 2 hours. CFpPS reactions contained prenyltransferase enriched extract and the corresponding prenyl-lipid donor. Other additives such as nanodiscs (see Supplementary methods for preparation of nanodiscs) and detergents were added at respectively indicated concentrations to the cell-free reactions.

#### Supported Lipid Bilayers (SLB) Formation

Preparation of glass coverslips (Menzel #1.5, 24x24 mm) and reaction chambers were performed according to previously protocols^61, 69^ and are in detail described in the Supplementary Methods. SLBs were formed through fusion of small unilamellar vesicles (SUVs) on the preformed reaction reservoir. SUVs were prepared as following: 80 mol% DOPC (Avanti Polar Lipids, Inc.), 19.95 mol% DOPS (Avanti Polar Lipids, Inc.) and 0.05 mol% ATTO488-DOPE (ATTO-TEC GmbH) were dissolved, mixed in chloroform, dried under a gentle stream of nitrogen, and transferred to a vacuum chamber for 1 hour. Then, the dried lipid film was rehydrated in SLB buffer A (50 mM Tris-HCl pH 7.5, 150 mM KCl, 5 mM MgCl_2_) to reach a final lipid concentration of 4 mg mL^-1^. Resulting samples were further vortexed and sonicated (bath sonicator, Branson) until they appeared clear. SLBs were prepared according to previous published protocols^70^. Briefly, SUVs (4 mg mL^-1^) were diluted with 130 μL SLB buffer A and 75 μL of the suspension were transferred to the preformed reaction chambers and incubated on a heat block at 37 °C for 1 min. 150 μL SLB buffer A were added into the chamber and incubated for further 2 min. The chamber was washed with 2 mL SLB buffer B (SLB buffer A without MgCl_2_) prior to a buffer-exchange to either prenylation buffer with 0.4% (w/v%) BSA (Sigma) for *in vitro* prenylation reactions or S30C buffer with BSA for CFpPS reactions, leaving 100 μL buffer inside the chamber to prevent drying of the formed SLBs. All used buffers should be pre-warmed to avoid temperature fluctuations.

### Reconstitution of <u>CAAX-protein</u> membrane targeting and extraction processes

CFpPS reactions were composed as indicated above, using corresponding plasmids without the GGPP substrate. 95 μL of the reaction mixtures were then transferred onto the SLB reservoir and the chambers were set-up on the confocal microscope equipped with a temperature controlling system (ibidi heating system, universal fit chamber). Reactions were then started by adding 5 μL GGPP to reach a final concentration of 10 μM. After the targeting process was finished, the SLBs were washed with prenylation buffer and RhoGDI was added to reach final concentration of 100 nM and 200 nM. Control experiments using purified mCherry-Cdc42 were performed accordingly, except that the CFpPS reaction was replaced by the *in vitro* prenylation reaction. Final reaction mixtures in SLB reservoirs contained: 1 μM mCherry-Cdc42, 250 nM GGTase-I, 10 μM GGPP, 0.4 % (w/v%) BSA in prenylation buffer. After the targeting process, the SLB was washed again with prenylation buffer to remove excess protein in solution and RhoGDI was added.

#### Microscopy and image analysis

Imaging was performed on a Zeiss LSM780/LSM800 confocal laser scanning microscope, using a Zeiss C-Apochromat 40X/1.20 water-immersion objective (Carl Zeiss AG, Oberkochen, Germany). A built-in definite focus was applied for imaging the time-series experiments. ATTO-488 (membrane-dye) was excited using the 488 nm laser, mCherry fusion proteins were excited using the 594 nm laser. For multicolor imaging, images for each channel were acquired sequentially to prevent bleed- through. The resolution was set up to 512x512 pixels. To visualize the membrane localization, z-stack images (perpendicular to the plane of the SLB) were obtained with time intervals of 45 s. For control experiments using purified proteins, the temperature was kept constant at 25 °C while for CFpPS reaction on SLBs the temperature was maintained at 30 ^°^C.

For analysis of membrane targeting, time series of z-stacks were processed using a custom Fiji macro (see Supplementary Methods). At each time point, the macro selected the membrane slice in the z-stack as the one with maximum mean intensity in the membrane channel, and compiled the mean intensity of the corresponding mCherry channel into a new file. The resulting fluorescence intensities were then plotted over time. The average intensity of the last slice into the solution was used to represent the intensity from the solution during the membrane targeting process. For membrane extraction, the intensities were calculated as the mean intensities of the brightest slice at each time point without further calibration. Three individual replicates were performed per experimental condition.

#### Reporting summary

Further information on research design is available in the Nature Research Reporting Summary linked to this article.

## Data availability

The datasets generated during and/or analyzed during the current study are available from the corresponding author on reasonable request.

## Acknowledgments

We would like to thank Prof. Dr. A. Itzen (Technical University of Munich) for the FTase and GGTase-I constructs and helpful discussions about their preparation. We thank Dr. Sabine Suppmann and the protein production core facility (Max Planck Institute of Biochemistry) for help with protein production and extract fermentation. We would like to thank M. Schaper for her help with plasmid preparations, Dr. K. Nakel and K. Anderson for their help in terms of protein production and purification, S. Bauer for lipid preparation assistance and H. Jia for the discussions regarding imaging processing. L.K., S. and P.S. have been supported by the MaxSynBio consortium which is jointly funded by the Federal Ministry of Education and Research of Germany and the Max Planck Society. T.H. and P.S. acknowledge funding through the Deutsche Forschungsgemeinschaft (DFG, German Research Foundation) - Project-ID 201269156 – SFB 1032 (A09).

## Author Contributions

L. K., S., T.H., and P. S. contributed to the experimental design and wrote the paper. L.K., T.H and S performed experiments and data analysis.

## Supplementary Information

### Supplementary Figures

**Supplementary Figure 1.**
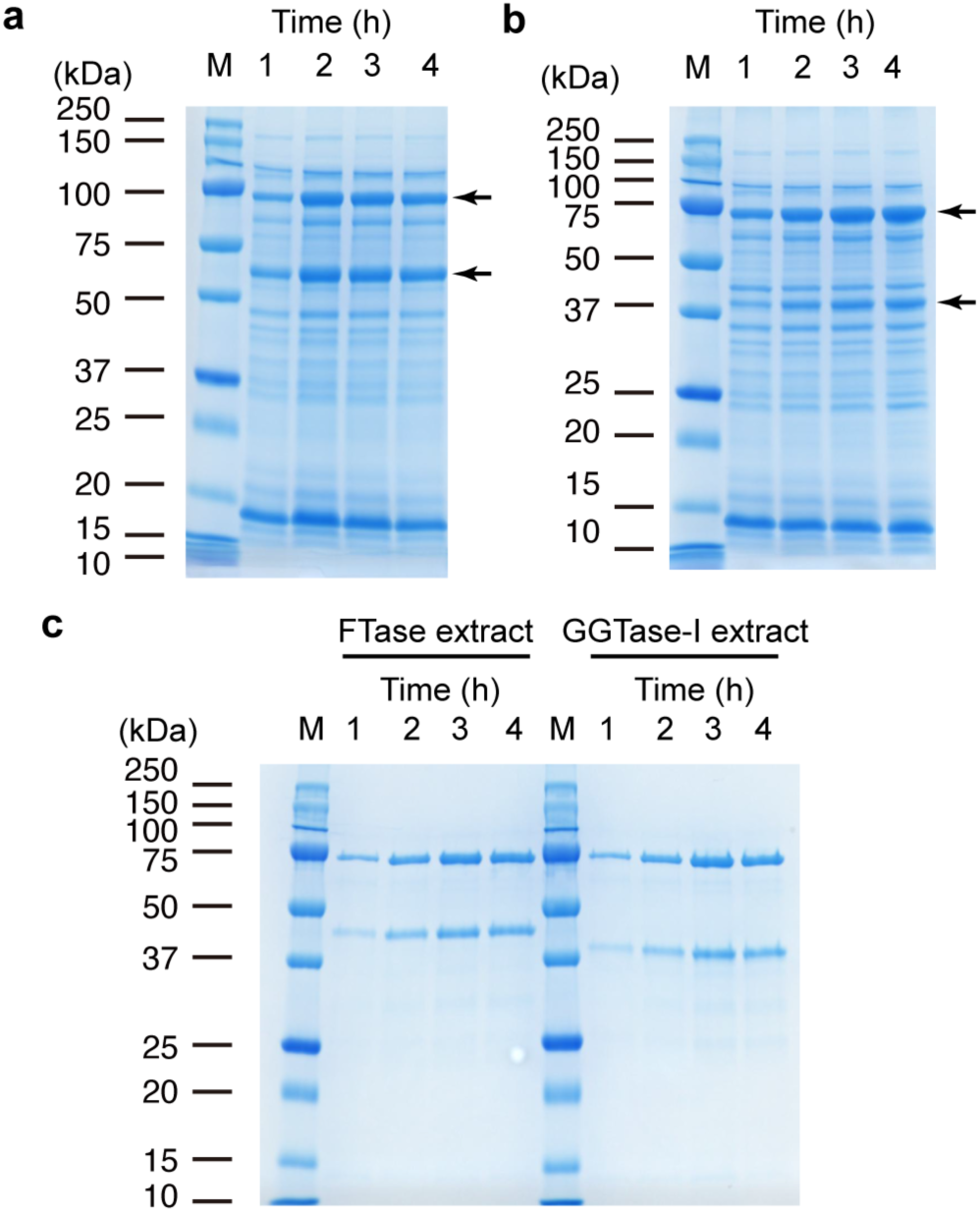
**Screening of induction time for the preparation of prenyltransferase-enriched extracts**. Coomassie-stained SDS-PAGE gels of clarified FTase **(a)** and GGTase-I **(b)** enriched cell lysates for varying time-points (1, 2, 3 or 4 h) after induction of expression. **(c)** Coomassie-stained gel of the IMAC-purified prenyltransferase-enriched lysates after induction of expression. Numbers above each lane represent the induction time in hours. Black arrows highlight the band corresponding to the *α* and *β* subunits of the prenyltransferase.

**Supplementary Figure 2.**
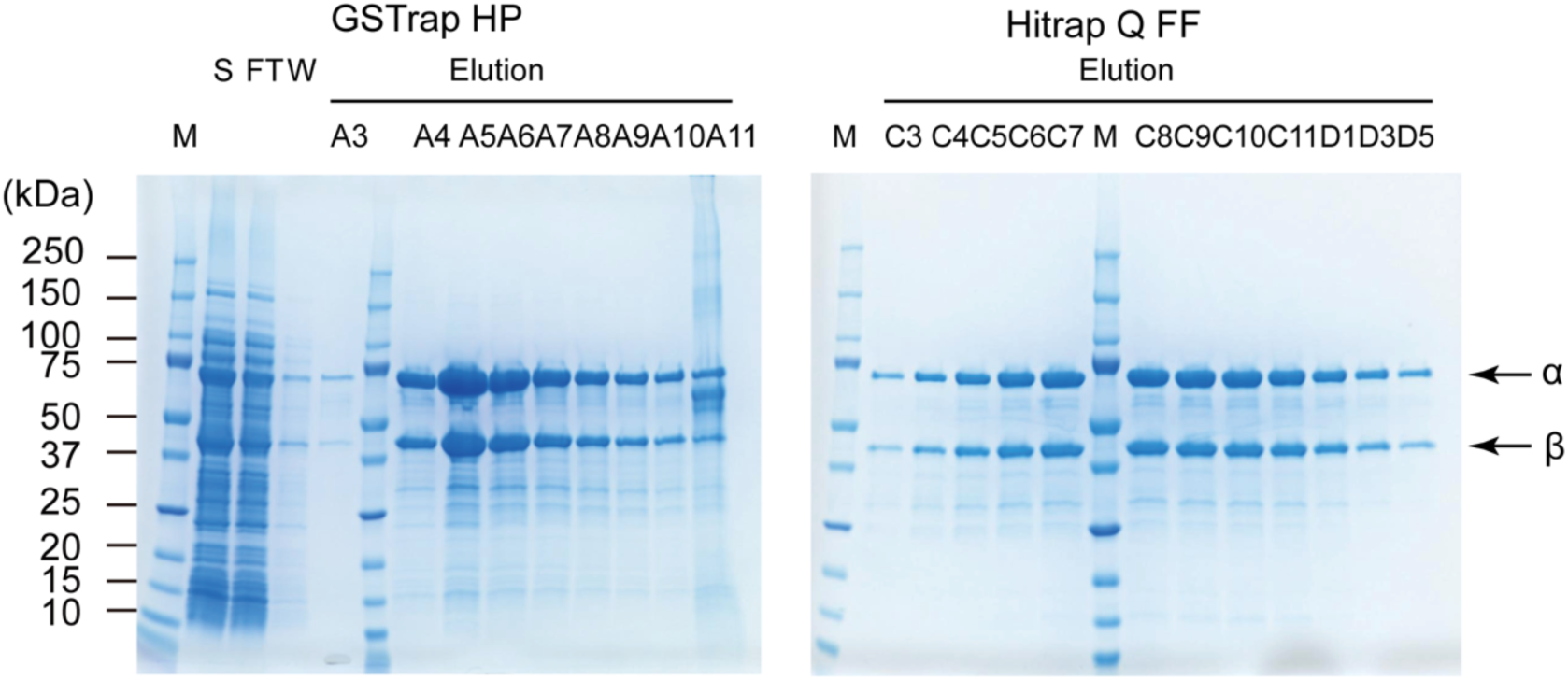
Purification of the prenyltransferase FTase. Coomassie- stained gels of collected fractions from the purification via a GSTrap HP column (left panel), followed by a Hitrap Q FF column (right panel). Lanes are titled with the fraction number obtained during elution. S indicates the supernatant, FT the flow-through, W the washing step and M denotes the protein marker. Black arrows indicate the bands that correspond to the *α* and *β* subunit of the FTase.

**Supplementary Figure 3.**
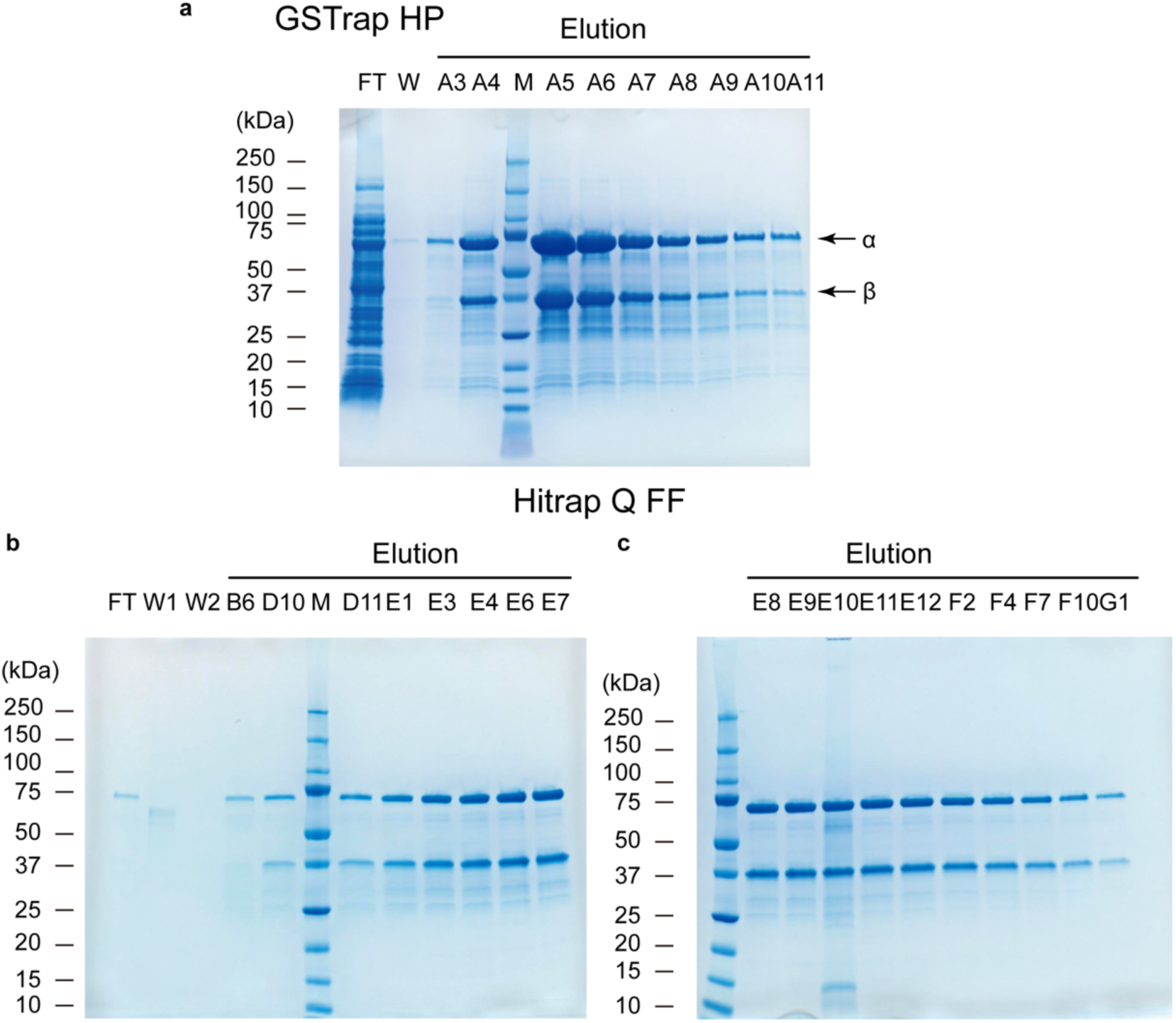
Purification of the prenyltransferase GGTase-I. Coomassie- stained gels of fractions collected from the purification via a GSTrap HP column **(a)**, followed by a Hitrap Q FF column **(b-c)**. Lanes are titled with the fraction number obtained during elution. FT indicates the flow-through and W, W1 and W2 the washing steps. M denotes the protein marker in all gels. Black arrows indicate the bands corresponding to the *α* and *β* subunit of the GGTase-I.

**Supplementary Figure 4.**
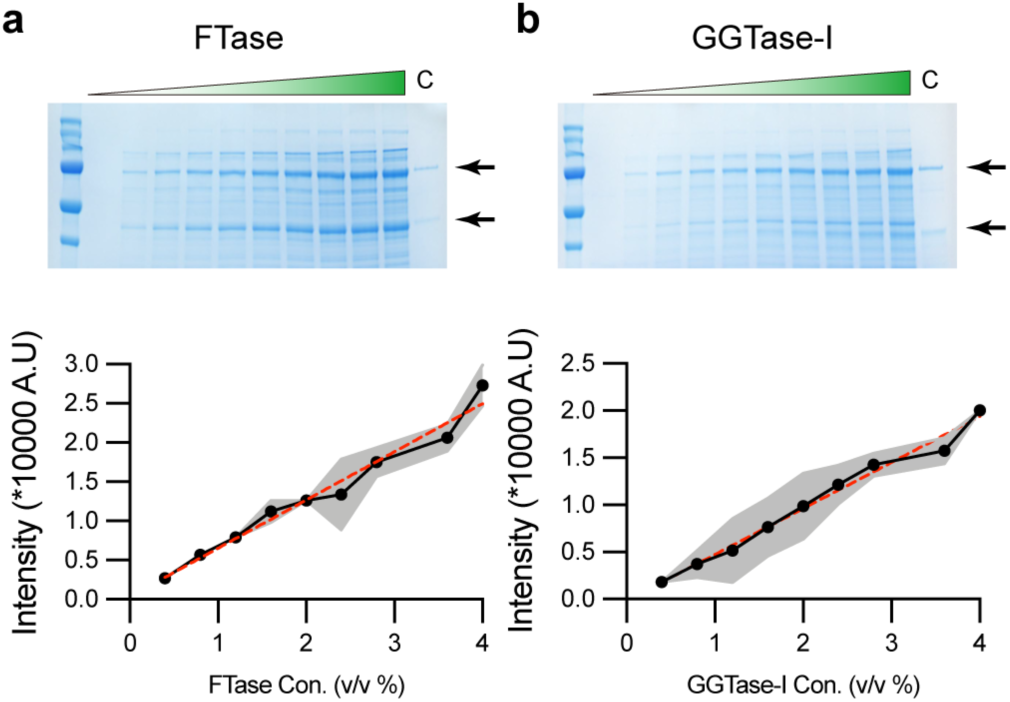
Crude estimation of the prenyltransferase concentration in enriched extracts. Coomassie-stained SDS-PAGE gels (upper row) of the serial dilution of either Ftase- **(a)** or GGTase-I-enriched **(b)** extract. A control sample (lane C) corresponding to the respective purified prenylatransferase and with a known concentration is shown for comparison (0.4 uM FTase, 2 uM GGTase-I). Lower panel: Estimation of the prenylatransfease concentration in the enriched extracts through colorimetry density estimation. Black arrows indicate the bands corresponding to the *α* and *β* subunit of the prenyltransferases.

**Supplementary Figure 5.**
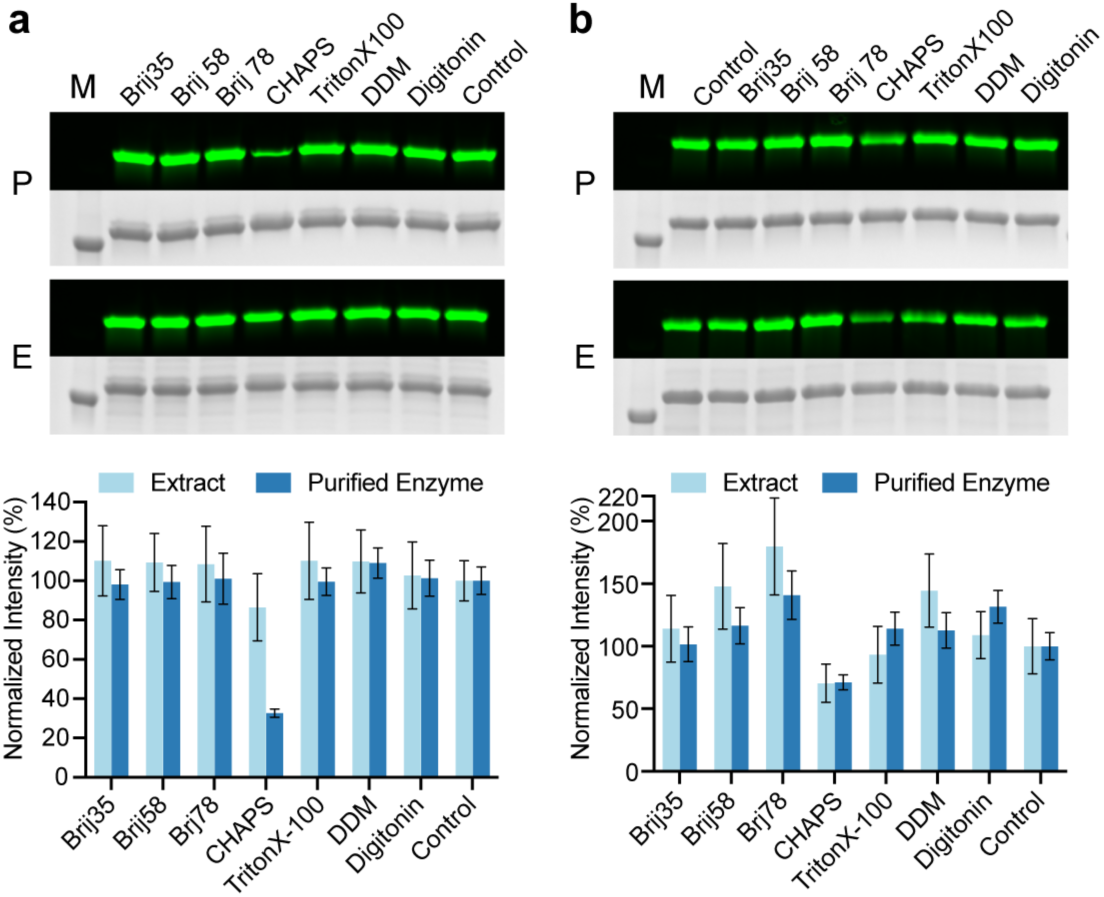
Detergent compatibility for *in vitro* prenylation reactions. Detergent compatibility for farnesylation **(a)** and for geranylgeranylation **(b)**. P denotes purified prenyltransferase, E denotes prenyltransferase-enriched extract and M denotes protein marker. In-gel fluorescence of NBD-labeled isoprenoid group was visualized for each modified protein band. The intensity of each fluorescent band was calibrated by densitometry from the corresponding Coomassie-stained gel images and normalized according to the control without any detergent. Bar graphs display mean values and standard deviations are shown, n = 3 independent replicates.

**Supplementary Figure 6.**
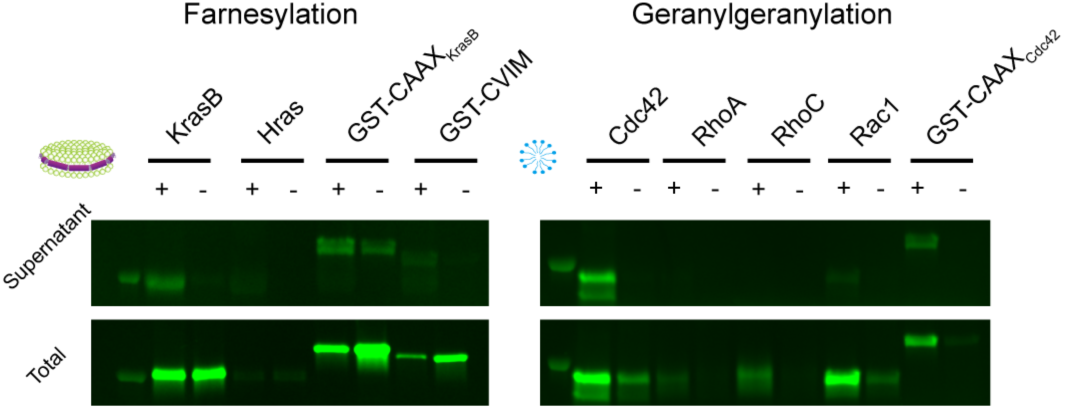
In-gel fluorescence analysis of the expression and solubilization of representative proteins. NBD-labeled isoprenoid group was used as lipid donor and visualized by in-gel fluorescence. Proteins correspond to those listed in Table 1 of the main manuscript. Nanodiscs (left panel) and detergents (right panel) were used for the farnesylation and geranylgerynlation of the exemplary proteins, respectively.

**Supplementary Figure 7.**
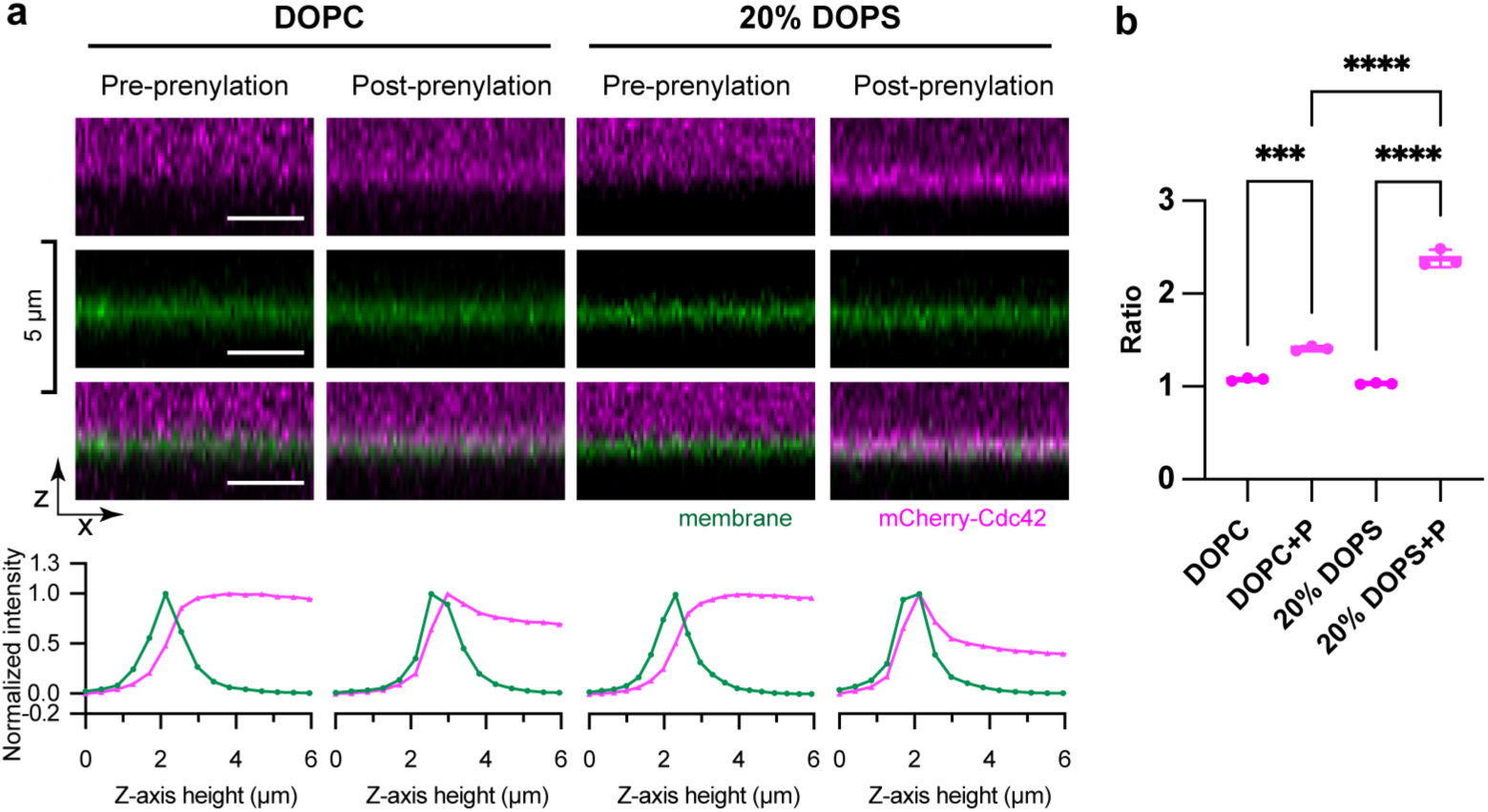
Charge-dependent targeting of mCherry-Cdc42 to supported lipid bilayers before and after prenylation. **(a)** Orthogonal views (upper panel) of mCherry-Cdc42 binding to neutral (100 % DOPC) or negatively charged (20% DOPS, 80% DOPC) supported lipid bilayers (SLB) before and after the addition of the lipid donor GGPP, membrane channel was visualized by the addition of 0.05 mol% ATTO488- DOPE. Lower panel: Normalized intensities of the respective Z-stacks for each experimental condition. **(b)** Ratio of the mCherry-Cdc42 intensity averaged over the Z-slice corresponding to the membrane and averaged over the slice of the stack furthest from the membrane (in solution). Abbreviations DOPC and 20 % DOPS stand for supported lipid bilayers containing only neural lipids or 20 % of negatively charged ones, respectively. The shortcut ‘+P’ indicates the presence of the lipid donor GGPP and hence a prenylated protein state. Statistical analysis of the determined ratios was performed by a one-way ANOVA followed by multiple comparisons. P values of 0.0001 are highlighted by *** while **** indicate P values of <0.0001, n = 3 independent replicates.

**Supplementary Figure 8.**
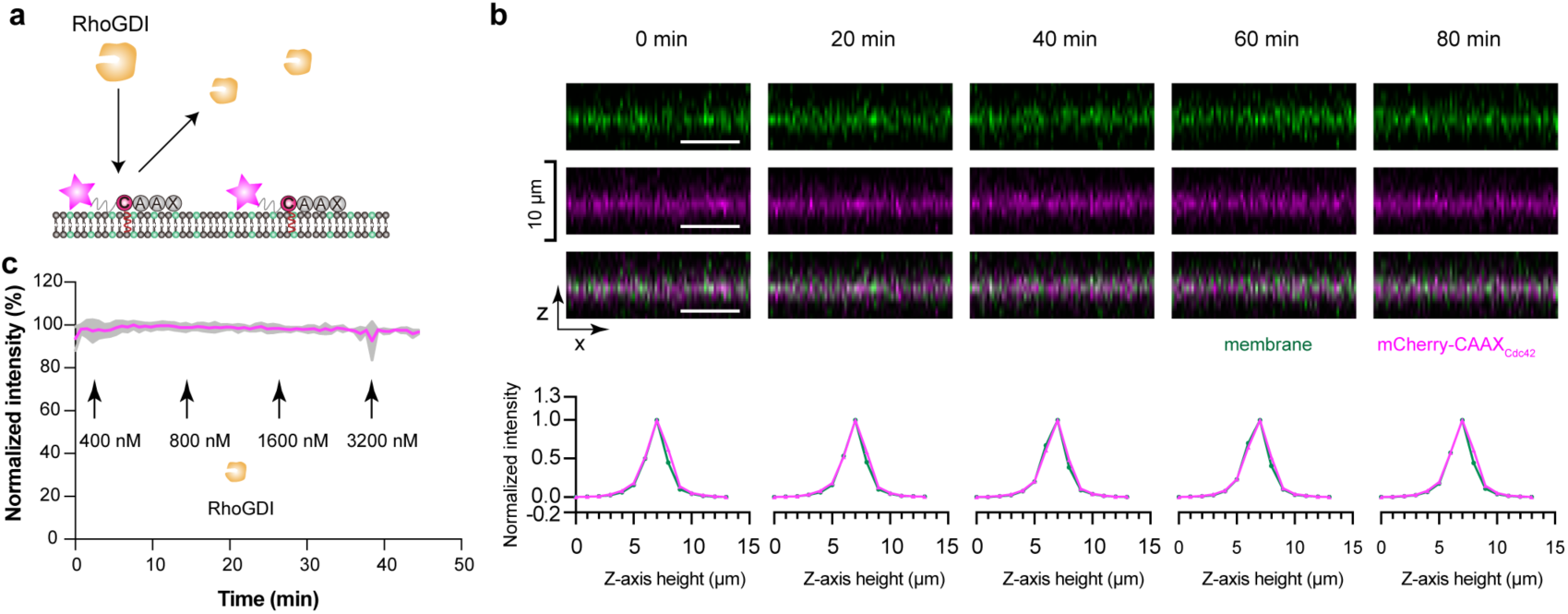
RhoGDI fails to extract mCherry-CAAX_Cdc42_ from the membrane. **(a)** Schematic illustration showing that unlike full-length Cdc42 (Fig. 4), RhoGDI cannot bind to mCherry-CAAX_Cdc42_ and extract it from the membrane. mCherry- CAAX_Cdc42_ lacks the interface required for RhoGDI interaction of Cdc42. **(b)** Orthogonal view of representative fluorescent images (upper panel) of membrane-localized mCherry- CAAX_Cdc42_ in the presence of RhoGDI. Lower panel, normalized fluorescence intensities of the corresponding images from upper panel. Images correspond to time since the start of imaging of an SLB (80% DOPC, 19.95% DOPS, 0.05% Atto-488 PE) loaded with prenylated mCherry-CAAX_Cdc42._ RhoGDI was added in increments at times observed in panel **c**. Membrane channel (green), mCherry-construct channel (magenta) and of the merged channels are displayed below. Scale bar: 10 µm. **(c)** Normalized average intensity of the mCherry-CAAX_Cdc42_ signal in the presence of RhoGDI. Standard deviations are indicated in gray shades (n=3) and a representative dataset of 3 independent replicates is shown. Intensity values were obtained by averaging over the slice that corresponds to the membrane and then normalized by setting the maximum and minimum intensities recorded during each experiment as 0 and 1, respectively. The indicated RhoGDI concentrations represent freshly added protein into the chamber at each time point besides the pre- existing RhoGDI.

**Supplementary Figure 9.**
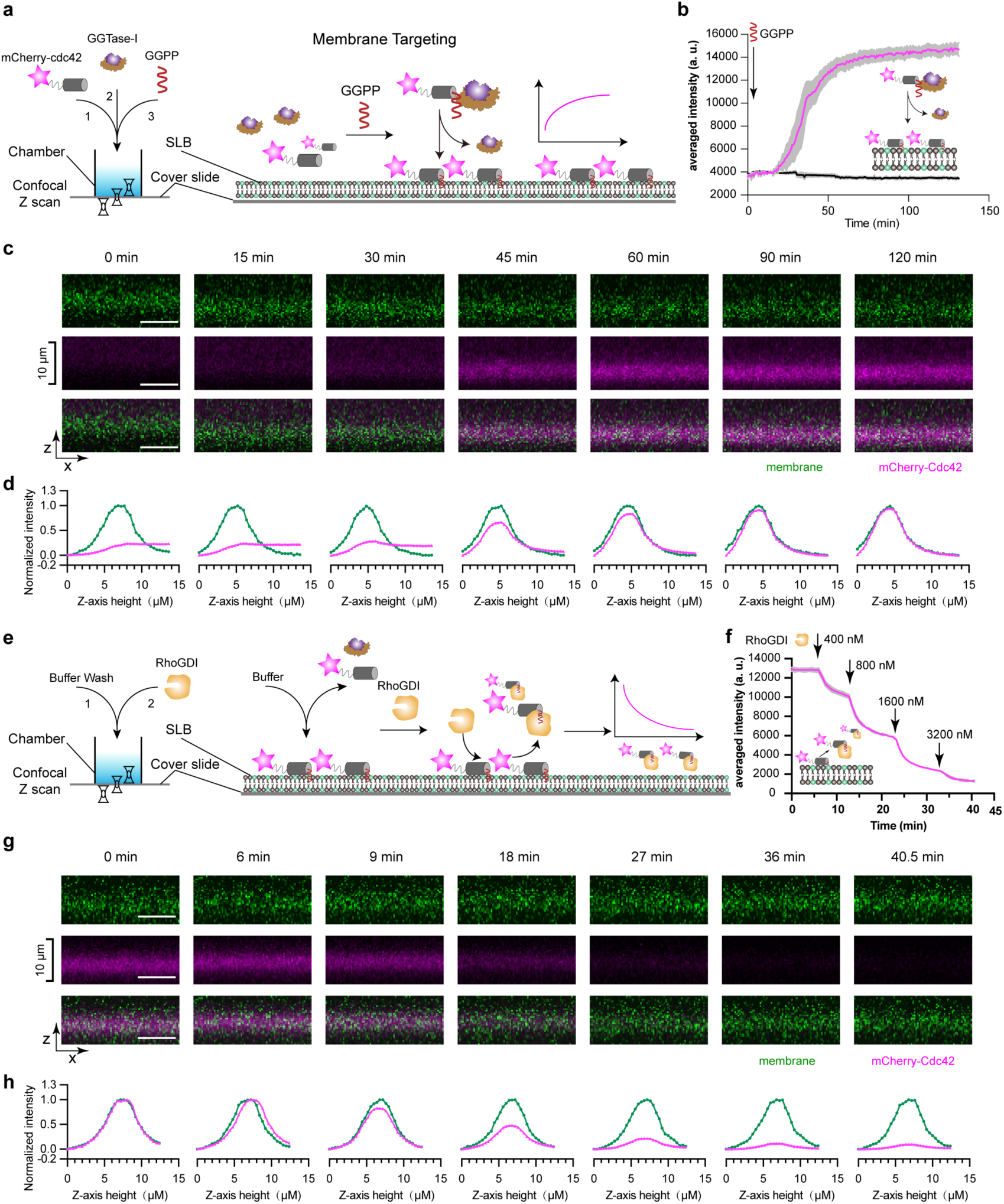
Reconstitution of the membrane targeting and extraction process of mCherry-Cdc42 using purified proteins. Schematic illustrations of membrane targeting **(a)** and extraction **(e)** processes reconstituted on supported lipid bilayers and imaged via a confocal laser scanning microscope. **(b, f)** Average intensities of mCherry-Cdc42 on the SLB membrane during targeting **(b)** and during RhoGDI- dependent membrane extraction **(f).** The SLB composition was 80% DOPC, 19.95% DOPS, 0.05% atto-488 PE. Standard deviations are shown in grey and the displayed graphs are a representative dataset of the 3 independent replicates. **(c, g)** Representative fluorescence images of the membrane targeting **(c)** and extraction process **(g)** of mCherry- Cdc42 (upper panel) in the x-y view. The corresponding orthogonal views of the membrane channel (green), the mCherry channel (magenta) and of the merged channels are displayed below. Scale bar: 10 µm. **(d, h)** Normalized intensities of the corresponding images displayed in panel (c) and (g), respectively. The membrane and mCherry channel intensities were normalized to the maximal (normalized to 1.0) and minimal intensity values (normalized to 0) from the z-stack during the targeting and extraction process.

## Supplementary Material and Methods

**Supplemental Table 1.** List of protein constructs used in this study

| Protein name | Vector | Source |
| --- | --- | --- |
| FTase $\alpha$ | pGEXTEV-FTase-α | 1 |
| FTase $\beta$ | pET28-FTase-β | 1 |
| GGTase-I $\alpha$ | pGEXTEV-GGTase-I-α | 1 |
| GGTase-I $\beta$ | pET28-GGTase-I-β | 1 |
| GST-CAAX<sub>KrasB</sub> | pGEX-RG-KrasB-C1-15 | This study |
| GST-CAAX<sub>Cdc42</sub> | pGEX-RG-hCdc42-C1-9 | This study |
| RhoGDI | pGEX-3C-RhoGDI | This study |
| mCherry-CAAX<sub>Cdc42</sub> | pCoofy1a-mCherry-RG-hCdc42-C1-9 | This study |
| RhoA | pIVEX2.3d-RhoA | This study |
| RhoC | pIVEX2.3d-RhoC | This study |
| Rac1 | pIVEX2.3d-Rac1 | This study |
| KrasB | pIVEX2.3d-KrasB | This study |
| HRas | pIVEX2.3d-HRas | This study |
| Cdc42 | pIVEX2.3d-Cdc42 | This study |
| mCherry-Cdc42 | pCoofy1a-mCherry-Cdc42 | This study |
| MSP1E3D1 | pET28a-MSP1E3D1 | 2 |

**Record of the complete sequences of genes and proteins that have been used or created in this study.**

### FTase *α*

Protein sequence MSPILGYWKIKGLVQPTRLLLEYLEEKYEEHLYERDEGDKWRNKKFELGLEFPNLPYYID GDVKLTQSMAIIRYIADKHNMLGGCPKERAEISMLEGAVLDIRYGVSRIAYSKDFETLKVD FLNKLPEMLKMFEDRLCHKTYLNGDHVTHPDFMLYDALDVVLYMDPMCLDAFPKLVCFK KRIEAIPQIDKYLKSSKYIAWPLQGWQATFGGGDHPPTNRWVSMADENLYFQGHMAAT EGVGESAPGGEPGQPEQPPPPPPPPPAQQPQEEEMAAEAGEAAASPMDDGFLSLDSP TYVLYRDRAEWADIDPVPQNDGPSPVVQIIYSEKFRDVYDYFRAVLQRDERSERAFKLT RDAIELNAANYTVWHFRRVLLRSLQKDLQEEMNYITAIIEEQPKNYQVWHHRRVLVEWL KDPSQELEFIADILNQDAKNYHAWQHRQWVIQEFRLWDNELQYVDQLLKEDVRNNSVW NQRHFVISNTTGYSDRAVLEREVQYTLEMIKLVPHNESAWNYLKGILQDRGLSRYPNLLN QLLDLQPSHSSPYLIAFLVDIYEDMLENQCDNKEDILNKALELCEILAKEKDTIRKEYWRYI GRSLQSKHSRESDIPASV*

#### DNA sequence

ATGTCCCCTATACTAGGTTATTGGAAAATTAAGGGCCTTGTGCAACCCACTCGACTTC TTTTGGAATATCTTGAAGAAAAATATGAAGAGCATTTGTATGAGCGCGATGAAGGTGA TAAATGGCGAAACAAAAAGTTTGAATTGGGTTTGGAGTTTCCCAATCTTCCTTATTATA TTGATGGTGATGTTAAATTAACACAGTCTATGGCCATCATACGTTATATAGCTGACAAG CACAACATGTTGGGTGGTTGTCCAAAAGAGCGTGCAGAGATTTCAATGCTTGAAGGA GCGGTTTTGGATATTAGATACGGTGTTTCGAGAATTGCATATAGTAAAGACTTTGAAAC TCTCAAAGTTGATTTTCTTAACAAGCTACCTGAAATGCTGAAAATGTTCGAAGATCGTT TATGTCATAAAACATATTTAAATGGTGATCATGTAACCCATCCTGACTTCATGTTGTATG ACGCTCTTGATGTTGTTTTATACATGGATCCAATGTGCCTGGATGCGTTCCCAAAATTA GTTTGTTTTAAAAAACGTATTGAAGCTATCCCACAAATTGATAAGTACTTGAAATCCAG CAAGTATATAGCATGGCCTTTGCAGGGCTGGCAAGCCACGTTTGGTGGTGGCGACC ATCCTCCAACTAATCGATGGGTATCCATGGCTGATGAGAATCTTTATTTTCAGGGCCAT ATGGCGGCCACTGAGGGGGTCGGGGAATCTGCGCCAGGCGGTGAGCCGGGACAG CCAGAGCAGCCGCCGCCCCCGCCTCCTCCGCCGCCAGCACAGCAGCCGCAGGAA GAAGAGATGGCGGCCGAGGCCGGGGAAGCAGCGGCGTCCCCTATGGACGACGGG TTTCTGAGCCTGGACTCGCCCACCTATGTCTTGTACAGGGACAGGGCAGAGTGGGC TGACATAGACCCAGTGCCCCAGAATGATGGCCCCAGTCCAGTGGTCCAGATCATCTA CAGTGAAAAGTTTAGAGACGTCTATGATTACTTCCGAGCTGTTCTGCAGCGCGATGA AAGAAGCGAACGAGCCTTTAAGCTCACTCGAGATGCTATTGAGTTAAACGCAGCCAA CTATACGGTGTGGCATTTTCGGAGAGTTCTCTTGAGGTCGCTTCAGAAGGATCTGCA AGAAGAAATGAACTACATCACTGCAATAATTGAGGAACAGCCCAAAAACTATCAAGTT TGGCACCATAGGAGAGTATTAGTGGAGTGGCTGAAAGATCCTTCTCAAGAGCTCGAG TTCATCGCCGATATCCTTAATCAGGATGCAAAGAATTACCATGCCTGGCAGCATCGAC AGTGGGTCATTCAGGAGTTTCGACTTTGGGATAATGAGCTGCAGTATGTGGACCAGC TTCTCAAAGAGGATGTGAGAAATAACTCTGTGTGGAACCAAAGACACTTCGTCATTTC TAATACCACTGGCTACAGTGATCGCGCTGTGTTGGAGAGAGAAGTCCAATATACTCTG GAAATGATCAAATTAGTGCCACACAATGAGAGTGCGTGGAACTACTTGAAAGGGATTT TGCAGGACCGTGGTCTTTCCAGATACCCTAATCTATTAAACCAGTTGCTTGATTTACAA CCAAGTCACAGCTCCCCCTACCTAATTGCCTTTCTTGTGGATATCTATGAAGACATGC TGGAAAACCAGTGTGACAACAAGGAGGACATTCTTAATAAAGCACTAGAGTTATGTGA GATTCTAGCTAAAGAAAAGGACACTATAAGAAAGGAATATTGGAGATATATTGGACGGT CCCTCCAGAGTAAACACAGCAGAGAAAGTGACATACCGGCGAGTGTATAG

### FTase *β*

Protein sequence MASSSSFTYYCPPSSSPVWSEPLYSLRPEHARERLQDDSVETVTSIEQAKVEEKIQEVF SSYKFNHLVPRLVLQREKHFHYLKRGLRQLTDAYECLDASRPWLCYWILHSLELLDEPIP QIVATDVCQFLELCQSPDGGFGGGPGQYPHLAPTYAAVNALCIIGTEEAYNVINREKLLQ YLYSLKQPDGSFLMHVGGEVDVRSAYCAASVASLTNIITPDLFEGTAEWIARCQNWEGGI GGVPGMEAHGGYTFCGLAALVILKKERSLNLKSLLQWVTSRQMRFEGGFQGRCNKLVD GCYSFWQAGLLPLLHRALHAQGDPALSMSHWMFHQQALQEYILMCCQCPAGGLLDKP GKSRDFYHTCYCLSGLSIAQHFGSGAMLHDVVMGVPENVLQPTHPVYNIGPDKVIQATT HFLQKPVPGFEECEDAVTSDPATD*

#### DNA sequence

ATGGCTTCTTCGAGTTCCTTCACCTATTATTGTCCTCCATCTTCTTCCCCTGTTTGGTC AGAACCGCTGTATAGTCTGAGACCTGAGCACGCGCGGGAGCGGTTGCAAGACGACT CAGTGGAAACAGTCACGTCCATAGAACAGGCCAAAGTAGAAGAAAAGATCCAGGAG GTCTTCAGTTCTTACAAGTTTAACCACCTCGTACCAAGGCTCGTTCTGCAGAGGGAG AAGCACTTCCATTATCTGAAAAGAGGCCTTCGACAACTGACAGATGCCTATGAGTGT CTGGATGCCAGCCGCCCCTGGCTCTGCTACTGGATCCTGCACAGCTTGGAGCTCCT CGACGAACCCATCCCCCAAATAGTGGCTACAGATGTGTGTCAGTTCTTGGAGCTGTG TCAGAGTCCAGACGGTGGCTTTGGAGGGGGCCCTGGTCAGTACCCACACCTCGCT CCCACGTATGCAGCTGTCAACGCGCTATGCATCATTGGCACGGAGGAAGCCTACAAC GTCATTAACAGAGAGAAGCTCCTTCAGTACTTGTACTCCCTAAAGCAACCGGATGGC TCTTTTCTCATGCACGTCGGAGGAGAGGTGGATGTAAGAAGTGCGTACTGTGCTGC CTCAGTAGCCTCTCTCACCAACATCATCACTCCTGACCTCTTCGAAGGCACTGCTGA ATGGATAGCAAGGTGCCAGAACTGGGAAGGCGGCATTGGCGGGGTGCCAGGGATG GAAGCCCACGGTGGCTACACCTTCTGTGGCTTGGCTGCGCTGGTGATCCTCAAGAA GGAACGTTCTTTGAACCTGAAGAGCTTGCTACAATGGGTGACAAGCCGGCAGATGC GGTTCGAAGGAGGATTTCAGGGCCGCTGCAACAAGCTGGTGGACGGCTGCTACTC CTTCTGGCAGGCAGGACTTCTGCCCCTGTTGCACCGGGCACTCCACGCTCAAGGT GACCCTGCCCTCAGCATGAGCCACTGGATGTTCCATCAGCAGGCGCTGCAGGAGTA CATCCTCATGTGCTGCCAGTGTCCGGCTGGGGGTCTCCTGGACAAACCTGGCAAGT CACGTGACTTCTACCATACTTGCTACTGCCTGAGCGGCCTGTCCATTGCCCAGCATT TTGGAAGTGGAGCCATGCTGCACGATGTGGTCATGGGTGTGCCTGAAAATGTTCTG CAGCCCACTCACCCTGTGTACAACATCGGACCTGATAAGGTGATCCAGGCCACCACA CACTTTCTGCAGAAGCCGGTCCCAGGCTTTGAGGAATGCGAAGATGCGGTGACCTC AGATCCTGCCACTGACTAG

### GGTase-I beta

Protein sequence MAATEDDRLAGSGEGERLDFLRDRHVRFFQRCLQVLPERYSSLETSRLTIAFFALSGLD MLDSLDVVNKDDIIEWIYSLQVLPTEDRSNLDRCGFRGSSYLGIPFNPSKNPGTAHPYDS GHIAMTYTGLSCLIILGDDLSRVDKEACLAGLRALQLEDGSFCAVPEGSENDMRFVYCA SCICYMLNNWSGMDMKKAISYIRRSMSYDNGLAQGAGLESHGGSTFCGIASLCLMGKL EEVFSEKELNRIKRWCIMRQQNGYHGRPNKPVDTCYSFWVGATLKLLKIFQYTNFEKNR NYILSTQDRLVGGFAKWPDSHPDALHAYFGICGLSLMEESGICKVHPALNVSTRTSERLR DLHQSWKTKDSKQCSDNVHISS*

#### DNA sequence

ATGGCGGCCACAGAGGATGACAGACTGGCGGGGAGCGGAGAAGGAGAACGGCTG GATTTCCTGCGGGACCGACACGTGCGGTTCTTCCAGCGCTGCCTCCAGGTCTTGCC GGAGCGGTATTCTTCGCTGGAGACCAGCAGGCTGACAATTGCATTTTTTGCACTCTC TGGGCTGGATATGTTGGACTCCTTGGATGTGGTGAACAAAGACGATATAATAGAGTG GATTTATTCCTTGCAGGTTCTTCCCACAGAAGACAGGTCAAATCTGGATCGCTGTGG TTTCCGAGGTTCTTCATATTTGGGTATTCCATTCAACCCATCAAAGAATCCAGGCACA GCTCATCCTTATGACAGTGGACACATAGCGATGACTTACACTGGTCTTTCCTGTTTAA TTATTCTTGGAGATGATTTAAGCCGAGTAGATAAAGAAGCTTGCTTAGCAGGCTTGAG AGCACTTCAGCTGGAAGATGGGAGCTTCTGTGCTGTTCCTGAAGGCAGTGAGAATG ACATGAGGTTTGTGTACTGTGCTTCCTGCATTTGCTATATGCTCAACAACTGGTCAGG CATGGATATGAAGAAAGCCATCAGCTACATTAGAAGAAGTATGTCCTATGACAATGGC CTGGCACAGGGGGCAGGACTTGAGTCTCATGGAGGATCCACCTTTTGTGGCATTGC

GTCACTGTGCCTGATGGGTAAACTGGAAGAAGTTTTTTCAGAGAAAGAACTGAACCG GATAAAGAGGTGGTGCATAATGAGGCAGCAGAACGGGTACCACGGAAGACCTAACA AGCCTGTCGACACCTGTTACTCTTTCTGGGTGGGAGCAACACTAAAGCTTTTGAAAA TTTTCCAGTACACTAACTTTGAGAAGAATAGGAATTACATCTTATCAACTCAGGATCGC CTTGTTGGGGGATTTGCTAAATGGCCAGACAGTCATCCAGATGCTTTGCATGCGTAC TTCGGGATCTGTGGCCTGTCACTAATGGAGGAGAGTGGAATTTGTAAAGTTCATCCT GCTCTGAATGTAAGCACACGAACTTCTGAGCGCCTCCGAGATCTCCATCAAAGCTGG AAGACCAAGGACTCTAAACAGTGCTCAGACAATGTCCATATTTCCAGTTGA

### GST-CAAX_KrasB_

Protein sequence MNTIHHHHHHNTSSNSMSPILGYWKIKGLVQPTRLLLEYLEEKYEEHLYERDEGDKWRN KKFELGLEFPNLPYYIDGDVKLTQSMAIIRYIADKHNMLGGCPKERAEISMLEGAVLDIRY GVSRIAYSKDFETLKVDFLNKLPEMLKMFEDRLCHKTYLNGDHVTHPDFMLYDALDVVL YMDPMCLDAFPKLVCFKKRIEAIPQIDKYLKSSKYIAWPLQGWQATFGGGDHPPGSAG*L AEAAAKEAAAKEAAAKEAAAKAAA***GKKKKKKSKTKCVIM**\*

#### DNA sequence

ATGAACACCATTCATCACCATCACCATCACAACACTAGTAGCAATTCCATGTCCCCTAT ACTAGGTTATTGGAAAATTAAGGGCCTTGTGCAACCCACTCGACTTCTTTTGGAATAT CTTGAAGAAAAATATGAAGAGCATTTGTATGAGCGCGATGAAGGTGATAAATGGCGAA ACAAAAAGTTTGAATTGGGTTTGGAGTTTCCCAATCTTCCTTATTATATTGATGGTGAT GTTAAATTAACACAGTCTATGGCCATCATACGTTATATAGCTGACAAGCACAACATGTT GGGTGGTTGTCCAAAAGAGCGTGCAGAGATTTCAATGCTTGAAGGAGCGGTTTTGG ATATTAGATACGGTGTTTCGAGAATTGCATATAGTAAAGACTTTGAAACTCTCAAAGTT GATTTTCTTAACAAGCTACCTGAAATGCTGAAAATGTTCGAAGATCGTTTATGTCATAA AACATATTTAAATGGTGATCATGTAACCCATCCTGACTTCATGTTGTATGACGCTCTTG ATGTTGTTTTATACATGGATCCAATGTGCCTGGATGCGTTCCCAAAATTAGTTTGTTTT AAAAAACGTATTGAAGCTATCCCACAAATTGATAAGTACTTGAAATCCAGCAAGTATAT AGCATGGCCTTTGCAGGGCTGGCAAGCCACGTTTGGTGGTGGCGACCATCCTCCAG GTTCTGCGGGTTTAGCAGAGGCGGCAGCTAAAGAAGCTGCTGCCAAAGAAGCTGCC GCGAAAGAAGCGGCTGCCAAGGCTGCGGCAGGCAAGAAAAAGAAAAAGAAAAGCA AGACAAAATGTGTAATCATGTGA

### GST-CAAX_Cdc42_

Protein sequence MNTIHHHHHHNTSSNSMSPILGYWKIKGLVQPTRLLLEYLEEKYEEHLYERDEGDKWRN KKFELGLEFPNLPYYIDGDVKLTQSMAIIRYIADKHNMLGGCPKERAEISMLEGAVLDIRY GVSRIAYSKDFETLKVDFLNKLPEMLKMFEDRLCHKTYLNGDHVTHPDFMLYDALDVVL YMDPMCLDAFPKLVCFKKRIEAIPQIDKYLKSSKYIAWPLQGWQATFGGGDHPPGSAG*L AEAAAKEAAAKEAAAKEAAAKAAA***KKSRRCVLL**\*

#### DNA sequence

ATGAACACCATTCATCACCATCACCATCACAACACTAGTAGCAATTCCATGTCCCCTAT ACTAGGTTATTGGAAAATTAAGGGCCTTGTGCAACCCACTCGACTTCTTTTGGAATAT CTTGAAGAAAAATATGAAGAGCATTTGTATGAGCGCGATGAAGGTGATAAATGGCGAA ACAAAAAGTTTGAATTGGGTTTGGAGTTTCCCAATCTTCCTTATTATATTGATGGTGAT GTTAAATTAACACAGTCTATGGCCATCATACGTTATATAGCTGACAAGCACAACATGTT GGGTGGTTGTCCAAAAGAGCGTGCAGAGATTTCAATGCTTGAAGGAGCGGTTTTGG ATATTAGATACGGTGTTTCGAGAATTGCATATAGTAAAGACTTTGAAACTCTCAAAGTT GATTTTCTTAACAAGCTACCTGAAATGCTGAAAATGTTCGAAGATCGTTTATGTCATAA AACATATTTAAATGGTGATCATGTAACCCATCCTGACTTCATGTTGTATGACGCTCTTG ATGTTGTTTTATACATGGATCCAATGTGCCTGGATGCGTTCCCAAAATTAGTTTGTTTT AAAAAACGTATTGAAGCTATCCCACAAATTGATAAGTACTTGAAATCCAGCAAGTATAT AGCATGGCCTTTGCAGGGCTGGCAAGCCACGTTTGGTGGTGGCGACCATCCTCCAG GTTCTGCGGGTTTAGCAGAGGCGGCAGCTAAAGAAGCTGCTGCCAAAGAAGCTGCC GCGAAAGAAGCGGCTGCCAAGGCTGCGGCAAAGAAATCCAGGCGGTGCGTTCTGC TGTGA

### GST-CVIM

Protein sequence MNTIHHHHHHNTSSNSMSPILGYWKIKGLVQPTRLLLEYLEEKYEEHLYERDEGDKWRN KKFELGLEFPNLPYYIDGDVKLTQSMAIIRYIADKHNMLGGCPKERAEISMLEGAVLDIRY GVSRIAYSKDFETLKVDFLNKLPEMLKMFEDRLCHKTYLNGDHVTHPDFMLYDALDVVL YMDPMCLDAFPKLVCFKKRIEAIPQIDKYLKSSKYIAWPLQGWQATFGGGDHPPGSAG*L AEAAAKEAAAKEAAAKEAAAKAAA***CVIM**\*

#### DNA sequence

ATGTCCCCTATACTAGGTTATTGGAAAATTAAGGGCCTTGTGCAACCCACTCGACTTC TTTTGGAATATCTTGAAGAAAAATATGAAGAGCATTTGTATGAGCGCGATGAAGGTGA TAAATGGCGAAACAAAAAGTTTGAATTGGGTTTGGAGTTTCCCAATCTTCCTTATTATA TTGATGGTGATGTTAAATTAACACAGTCTATGGCCATCATACGTTATATAGCTGACAAG CACAACATGTTGGGTGGTTGTCCAAAAGAGCGTGCAGAGATTTCAATGCTTGAAGGA GCGGTTTTGGATATTAGATACGGTGTTTCGAGAATTGCATATAGTAAAGACTTTGAAAC TCTCAAAGTTGATTTTCTTAACAAGCTACCTGAAATGCTGAAAATGTTCGAAGATCGTT TATGTCATAAAACATATTTAAATGGTGATCATGTAACCCATCCTGACTTCATGTTGTATG ACGCTCTTGATGTTGTTTTATACATGGATCCAATGTGCCTGGATGCGTTCCCAAAATTA GTTTGTTTTAAAAAACGTATTGAAGCTATCCCACAAATTGATAAGTACTTGAAATCCAG CAAGTATATAGCATGGCCTTTGCAGGGCTGGCAAGCCACGTTTGGTGGTGGCGACC ATCCTCCAGGTTCTGCGGGTTTAGCAGAGGCGGCAGCTAAAGAAGCTGCTGCCAAA GAAGCTGCCGCGAAAGAAGCGGCTGCCAAGGCTGCGGCATGTGTAATCATGTGA

### RhoGDI

Protein sequence MSPILGYWKIKGLVQPTRLLLEYLEEKYEEHLYERDEGDKWRNKKFELGLEFPNLPYYID GDVKLTQSMAIIRYIADKHNMLGGCPKERAEISMLEGAVLDIRYGVSRIAYSKDFETLKVD FLSKLPEMLKMFEDRLCHKTYLNGDHVTHPDFMLYDALDVVLYMDPMCLDAFPKLVCFK KRIEAIPQIDKYLKSSKYIAWPLQGWQATFGGGDHPPKSDLEVLFQGPLGSPEFMAEQE PTAEQLAQIAAENEEDEHSVNYKPPAQKSIQEIQELDKDDESLRKYKEALLGRVAVSADP NVPNVVVTGLTLVCSSAPGPLELDLTGDLESFKKQSFVLKEGVEYRIKISFRVNREIVSGM KYIQHTYRKGVKIDKTDYMVGSYGPRAEEYEFLTPVEEAPKGMLARGSYSIKSRFTDDD KTDHLSWEWNLTIKKDWKD*

#### DNA sequence

ATGTCCCCTATACTAGGTTATTGGAAAATTAAGGGCCTTGTGCAACCCACTCGACTTC TTTTGGAATATCTTGAAGAAAAATATGAAGAGCATTTGTATGAGCGCGATGAAGGTGA TAAATGGCGAAACAAAAAGTTTGAATTGGGTTTGGAGTTTCCCAATCTTCCTTATTATA TTGATGGTGATGTTAAATTAACACAGTCTATGGCCATCATACGTTATATAGCTGACAAG CACAACATGTTGGGTGGTTGTCCAAAAGAGCGTGCAGAGATTTCAATGCTTGAAGGA GCGGTTTTGGATATTAGATACGGTGTTTCGAGAATTGCATATAGTAAAGACTTTGAAAC TCTCAAAGTTGATTTTCTTAGCAAGCTACCTGAAATGCTGAAAATGTTCGAAGATCGT TTATGTCATAAAACATATTTAAATGGTGATCATGTAACCCATCCTGACTTCATGTTGTAT GACGCTCTTGATGTTGTTTTATACATGGACCCAATGTGCCTGGATGCGTTCCCAAAAT TAGTTTGTTTTAAAAAACGTATTGAAGCTATCCCACAAATTGATAAGTACTTGAAATCCA GCAAGTATATAGCATGGCCTTTGCAGGGCTGGCAAGCCACGTTTGGTGGTGGCGAC CATCCTCCAAAATCGGATCTGGAAGTTCTGTTCCAGGGGCCCCTGGGATCCCCGGA ATTCATGGCCGAGCAAGAACCGACTGCAGAACAACTTGCGCAAATTGCAGCGGAAA ACGAGGAAGATGAGCATAGCGTGAACTACAAACCACCAGCCCAGAAAAGCATTCAG GAAATTCAGGAGCTGGATAAAGACGATGAATCGCTGCGGAAATACAAAGAAGCCCTC TTAGGTCGTGTAGCGGTTTCAGCGGATCCGAATGTCCCGAATGTTGTGGTGACTGGC CTGACGTTGGTCTGCAGCAGTGCTCCTGGTCCGTTAGAGCTGGATCTGACGGGTGA TCTGGAATCGTTCAAGAAACAGAGCTTTGTCCTGAAAGAAGGGGTGGAATATCGCAT CAAAATCTCTTTTCGCGTAAATCGCGAAATTGTGTCTGGCATGAAATACATTCAGCAC ACCTATCGCAAAGGCGTGAAAATCGACAAAACGGACTATATGGTTGGATCGTATGGTC CTCGTGCGGAAGAGTATGAGTTCCTCACACCGGTTGAAGAAGCACCCAAAGGCATG CTTGCTCGTGGGTCCTACTCCATTAAGTCACGCTTTACCGACGATGACAAGACCGAT CATCTGAGTTGGGAATGGAACTTGACCATCAAGAAAGACTGGAAAGATTGA

### mCherry-CAAX_Cdc42_

Protein sequence MKHHHHHHHHHHSAGLEVLFQGPMVSKGEEDNMAIIKEFMRFKVHMEGSVNGHEFEIE GEGEGRPYEGTQTAKLKVTKGGPLPFAWDILSPQFMYGSKAYVKHPADIPDYLKLSFPE GFKWERVMNFEDGGVVTVTQDSSLQDGEFIYKVKLRGTNFPSDGPVMQKKTMGWEA SSERMYPEDGALKGEIKQRLKLKDGGHYDAEVKTTYKAKKPVQLPGAYNVNIKLDITSH NEDYTIVEQYERAEGRHSTGGMDELYK*LAEAAAKEAAAKEAAAKEAAAKAAA***KKSRRC VLL**\*

#### DNA sequence

ATGAAACATCACCATCACCATCACCATCATCACCATTCCGCGGGTCTGGAAGTTCTGT TCCAGGGGCCCATGGTGAGCAAGGGCGAGGAGGATAACATGGCCATCATCAAGGAG TTCATGCGCTTCAAGGTGCACATGGAGGGCTCCGTGAACGGCCACGAGTTCGAGAT CGAGGGCGAGGGCGAGGGCCGCCCCTACGAGGGCACCCAGACCGCCAAGCTGAA GGTGACCAAGGGTGGCCCCCTGCCCTTCGCCTGGGACATCCTGTCCCCTCAGTTCA TGTACGGCTCCAAGGCCTACGTGAAGCACCCCGCCGACATCCCCGACTACTTGAAG CTGTCCTTCCCCGAGGGCTTCAAGTGGGAGCGCGTGATGAACTTCGAGGACGGCG GCGTGGTGACCGTGACCCAGGACTCCTCCCTGCAGGACGGCGAGTTCATCTACAAG GTGAAGCTGCGCGGCACCAACTTCCCCTCCGACGGCCCCGTAATGCAGAAGAAGA CCATGGGCTGGGAGGCCTCCTCCGAGCGGATGTACCCCGAGGACGGCGCCCTGAA GGGCGAGATCAAGCAGAGGCTGAAGCTGAAGGACGGCGGCCACTACGACGCTGAG GTCAAGACCACCTACAAGGCCAAGAAGCCCGTGCAGCTGCCCGGCGCCTACAACG TCAACATCAAGTTGGACATCACCTCCCACAACGAGGACTACACCATCGTGGAACAGT ACGAACGCGCCGAGGGCCGCCACTCCACCGGCGGCATGGACGAGCTGTACAAGTT AGCAGAGGCGGCAGCTAAAGAAGCTGCTGCCAAAGAAGCTGCCGCGAAAGAAGCG GCTGCCAAGGCTGCGGCAAAGAAATCCAGGCGGTGCGTTCTGCTGTGA

### RhoA

Protein sequence MKHHHHHHHHHHSAGLEVLFQGPMAAIRKKLVIVGDGACGKTCLLIVFSKDQFPEVYVP TVFENYVADIEVDGKQVELALWDTAGQEDYDRLRPLSYPDTDVILMCFSIDSPDSLENIP EKWTPEVKHFCPNVPIILVGNKKDLRNDEHTRRELAKMKQEPVKPEEGRDMANRIGAF GYMECSAKTKDGVREVFEMATRAALQARRGKKKSGCLVL*

#### DNA sequence

ATGAAACATCACCATCACCATCACCATCATCACCATTCCGCGGGTCTGGAAGTTCTGT TCCAGGGGCCCATGGCGGCAATTCGCAAGAAACTGGTGATTGTAGGCGATGGTGCC TGTGGCAAAACCTGTCTGCTGATCGTGTTCTCCAAAGACCAGTTTCCTGAAGTGTAT GTTCCCACCGTATTCGAAAACTACGTAGCGGACATTGAGGTTGATGGCAAACAGGTG GAATTAGCCCTGTGGGATACCGCAGGGCAAGAAGATTATGACCGGTTGCGCCCGTTA AGCTATCCGGATACGGACGTCATCCTGATGTGCTTCAGCATCGATTCGCCAGATTCTC TCGAGAACATTCCGGAGAAATGGACACCAGAAGTCAAACACTTTTGCCCGAATGTTC CGATTATCCTGGTGGGCAATAAGAAAGACCTTCGCAACGATGAGCATACCCGTCGCG AACTGGCGAAAATGAAACAGGAACCTGTCAAACCGGAAGAGGGACGTGACATGGCC AATCGCATTGGTGCGTTTGGGTACATGGAATGCAGTGCGAAAACGAAAGATGGTGTT CGCGAAGTGTTTGAAATGGCCACTCGTGCTGCTCTCCAAGCACGTCGTGGTAAGAA GAAATCAGGCTGTTTGGTCCTGTGA

### RhoC

Protein sequence MKHHHHHHHHHHSAGLEVLFQGPMAAIRKKLVIVGDGACGKTCLLIVFSKDQFPEVYVP TVFENYIADIEVDGKQVELALWDTAGQEDYDRLRPLSYPDTDVILMCFSIDSPDSLENIPE KWTPEVKHFCPNVPIILVGNKKDLRQDEHTRRELAKMKQEPVRSEEGRDMANRISAFG YLECSAKTKEGVREVFEMATRAGLQVRKNKRRRGCPIL*

#### DNA sequence

ATGAAACATCACCATCACCATCACCATCATCACCATTCCGCGGGTCTGGAAGTTCTGT TCCAGGGGCCCATGGCAGCAATTCGCAAGAAACTGGTGATTGTCGGCGATGGTGCC TGTGGCAAAACCTGTCTGCTGATTGTGTTCTCCAAAGACCAGTTTCCTGAAGTGTAT GTGCCGACAGTTTTCGAGAACTACATTGCGGATATCGAAGTAGACGGGAAACAGGTG GAACTGGCGTTATGGGATACGGCTGGTCAGGAAGATTACGATCGCCTGCGTCCGTT GAGCTATCCGGATACCGATGTCATCCTCATGTGCTTCTCGATTGACAGCCCTGACAG TCTGGAAAATATCCCCGAGAAATGGACGCCAGAAGTCAAGCACTTTTGCCCGAATGT ACCGATCATCCTTGTAGGCAACAAGAAAGATCTGCGCCAAGACGAACATACCCGTCG CGAATTGGCGAAAATGAAACAAGAGCCAGTTCGGTCAGAAGAAGGTCGCGATATGG CTAATCGCATTTCGGCCTTTGGCTATCTGGAATGCTCTGCGAAAACCAAAGAAGGAG TTCGCGAGGTGTTTGAGATGGCCACTCGTGCAGGGTTACAGGTTCGCAAGAACAAA CGTCGTCGGGGTTGTCCGATTCTCTAA

### Rac1

Protein sequence MKHHHHHHHHHHSAGLEVLFQGPMQAIKCVVVGDGAVGKTCLLISYTTNAFPGEYIPTV FDNYSANVMVDGKPVNLGLWDTAGQEDYDRLRPLSYPQTDVFLICFSLVSPASFENVRA KWYPEVRHHCPNTPIILVGTKLDLRDDKDTIEKLKEKKLTPITYPQGLAMAKEIGAVKYLE CSALTQRGLKTVFDEAIRAVLCPPPVKKRKRKCLLL*

#### DNA sequence

ATGAAACATCACCATCACCATCACCATCATCACCATTCCGCGGGTCTGGAAGTTCTGT TCCAGGGGCCCATGCAGGCGATCAAATGCGTAGTTGTCGGCGATGGTGCCGTCGGA AAGACCTGTCTGCTGATTAGCTACACCACAAACGCGTTTCCTGGCGAATACATCCCA ACCGTTTTCGACAACTATTCGGCGAATGTGATGGTTGATGGGAAACCCGTGAATCTG GGCTTGTGGGATACTGCTGGTCAGGAGGATTACGATCGCTTACGCCCACTTAGCTAT CCACAGACAGACGTCTTTCTGATCTGCTTTTCACTGGTGTCTCCCGCTTCCTTCGAA AACGTACGCGCCAAATGGTATCCGGAAGTTCGTCACCATTGCCCGAATACGCCGATT ATCCTCGTAGGTACCAAACTCGATTTGCGCGATGACAAAGACACGATTGAGAAACTG AAGGAAAAGAAACTGACGCCGATTACCTATCCGCAAGGCTTAGCAATGGCCAAAGAG ATTGGTGCAGTCAAATATCTGGAATGCAGTGCACTGACTCAACGTGGGCTGAAAACC GTGTTCGATGAAGCGATTCGTGCGGTGTTATGTCCGCCTCCGGTGAAGAAACGGAA ACGCAAATGTTTGCTGCTTTGA

### KrasB

Protein sequence MKHHHHHHHHHHSAGLEVLFQGPMTEYKLVVVGAGGVGKSALTIQLIQNHFVDEYDPTI EDSYRKQVVIDGETCLLDILDTAGQEEYSAMRDQYMRTGEGFLCVFAINNTKSFEDIHHY REQIKRVKDSEDVPMVLVGNKCDLPSRTVDTKQAQDLARSYGIPFIETSAKTRQGVDDA FYTLVREIRKHKEKMSKDGKKKKKKSKTKCVIM*

#### DNA sequence

ATGAAACATCACCATCACCATCACCATCATCACCATTCCGCGGGTCTGGAAGTTCTGT TCCAGGGGCCCATGACCGAATACAAACTGGTCGTAGTGGGTGCTGGTGGTGTTGGC AAATCAGCGTTGACCATTCAGCTGATTCAGAATCACTTCGTGGATGAGTATGATCCGA CAATTGAGGATAGCTATCGCAAACAGGTAGTGATTGACGGCGAAACTTGCCTCTTAG ACATTCTGGATACCGCTGGGCAAGAAGAGTACTCTGCAATGCGCGATCAGTACATGC GTACTGGTGAAGGCTTTCTGTGTGTCTTTGCGATCAACAATACCAAGAGCTTCGAAG ATATCCATCATTATCGGGAACAGATCAAACGTGTGAAAGACTCGGAAGATGTGCCAAT GGTCCTTGTTGGCAACAAATGCGATCTGCCTAGTCGTACCGTTGACACGAAACAAGC CCAGGACTTAGCACGCAGTTATGGCATTCCGTTCATTGAAACATCCGCCAAAACCCG TCAAGGAGTTGATGATGCGTTTTATACGCTGGTACGCGAAATCCGCAAACACAAAGA GAAAATGTCGAAAGACGGGAAGAAGAAGAAAAAGAAAAGCAAAACGAAATGTGTGAT CATGTGA

### HRas

Protein sequence MKHHHHHHHHHHSAGLEVLFQGPMTEYKLVVVGAGGVGKSALTIQLIQNHFVDEYDPTI EDSYRKQVVIDGETCLLDILDTAGQEEYSAMRDQYMRTGEGFLCVFAINNTKSFEDIHQY REQIKRVKDSDDVPMVLVGNKCDLAARTVESRQAQDLARSYGIPYIETSAKTRQGVEDA FYTLVREIRQHKLRKLNPPDESGPGCMSCKCVLS*

#### DNA sequence

ATGAAACATCACCATCACCATCACCATCATCACCATTCCGCGGGTCTGGAAGTTCTGT TCCAGGGGCCCATGACCGAATACAAGCTGGTGGTCGTAGGGGCTGGAGGTGTTGG CAAAAGTGCGTTGACCATCCAGCTGATTCAGAACCACTTCGTGGATGAGTACGACCC CACCATTGAAGATAGCTATCGCAAACAGGTCGTGATTGACGGAGAGACTTGTTTGCT GGACATTCTGGATACAGCTGGCCAAGAAGAGTATAGCGCAATGCGCGATCAGTACAT GCGTACCGGTGAAGGCTTTCTGTGCGTGTTTGCGATCAACAACACGAAATCCTTTGA GGACATCCATCAGTATCGCGAACAGATCAAACGCGTCAAAGACAGCGATGATGTGCC GATGGTTCTGGTTGGGAATAAGTGCGATTTAGCCGCACGTACGGTTGAATCGCGTCA AGCGCAAGATCTTGCCCGTTCCTATGGCATTCCGTATATCGAAACCTCTGCCAAAACG CGTCAAGGTGTAGAAGATGCGTTCTACACTCTGGTACGCGAAATTCGGCAGCATAAA CTGCGCAAACTCAATCCGCCAGACGAAAGTGGTCCTGGCTGTATGTCGTGTAAGTG CGTGTTATCATGA

### Cdc42

Protein sequence MKHHHHHHHHHHSAGLEVLFQGPMQTIKCVVVGDGAVGKTCLLISYTTNKFPSEYVPT VFDNYAVTVMIGGEPYTLGLFDTAGQEDYDRLRPLSYPQTDVFLVCFSVVSPSSFENVK EKWVPEITHHCPKTPFLLVGTQIDLRDDPSTIEKLAKNKQKPITPETAEKLARDLKAVKYV ECSALTQKGLKNVFDEAILAALEPPEPKKSRRCVLL*

#### DNA sequence

ATGAAACATCACCATCACCATCACCATCATCACCATTCCGCGGGTCTGGAAGTTCTGT TCCAGGGGCCCATGCAGACCATCAAATGCGTAGTCGTGGGTGATGGAGCAGTTGGC AAAACGTGTCTGCTCATCTCCTATACCACCAACAAATTTCCCAGCGAATACGTGCCTA CGGTATTCGACAATTACGCGGTTACCGTCATGATTGGTGGTGAACCCTATACCTTGGG CCTGTTCGATACTGCAGGCCAAGAGGACTATGATCGCTTACGCCCTCTGTCGTATCC GCAAACCGACGTCTTTCTTGTGTGCTTTAGCGTTGTGTCTCCGAGTTCGTTCGAAAA CGTGAAAGAGAAATGGGTACCGGAAATTACGCACCATTGTCCGAAAACTCCGTTTCT GCTGGTTGGCACACAGATCGATCTGCGCGATGATCCAAGCACCATTGAGAAACTTGC CAAGAACAAACAGAAACCGATTACGCCAGAAACTGCGGAGAAATTAGCCCGTGATCT GAAAGCCGTCAAGTACGTGGAATGCTCAGCTTTGACACAGAAAGGGCTGAAGAATG TGTTTGACGAAGCGATTCTGGCTGCGTTAGAACCGCCAGAACCGAAGAAAAGTCGT CGGTGTGTTCTCCTGTGA

### mCherry-Cdc42

Protein sequence MKHHHHHHHHHHSAGLEVLFQGPMVSKGEEDNMAIIKEFMRFKVHMEGSVNGHEFEIE GEGEGRPYEGTQTAKLKVTKGGPLPFAWDILSPQFMYGSKAYVKHPADIPDYLKLSFPE GFKWERVMNFEDGGVVTVTQDSSLQDGEFIYKVKLRGTNFPSDGPVMQKKTMGWEA SSERMYPEDGALKGEIKQRLKLKDGGHYDAEVKTTYKAKKPVQLPGAYNVNIKLDITSH NEDYTIVEQYERAEGRHSTGGMDELYKEFMQTIKCVVVGDGAVGKTCLLISYTTNKFPS EYVPTVFDNYAVTVMIGGEPYTLGLFDTAGQEDYDRLRPLSYPQTDVFLVCFSVVSPSS FENVKEKWVPEITHHCPKTPFLLVGTQIDLRDDPSTIEKLAKNKQKPITPETAEKLARDLK AVKYVECSALTQKGLKNVFDEAILAALEPPEPKKSRRCVLL*

#### DNA sequence

ATGAAACATCACCATCACCATCACCATCATCACCATTCCGCGGGTCTGGAAGTTCTGT TCCAGGGGCCCATGGTGAGCAAGGGCGAGGAGGATAACATGGCCATCATCAAGGAG TTCATGCGCTTCAAGGTGCACATGGAGGGCTCCGTGAACGGCCACGAGTTCGAGAT CGAGGGCGAGGGCGAGGGCCGCCCCTACGAGGGCACCCAGACCGCCAAGCTGAA GGTGACCAAGGGTGGCCCCCTGCCCTTCGCCTGGGACATCCTGTCCCCTCAGTTCA TGTACGGCTCCAAGGCCTACGTGAAGCACCCCGCCGACATCCCCGACTACTTGAAG CTGTCCTTCCCCGAGGGCTTCAAGTGGGAGCGCGTGATGAACTTCGAGGACGGCG GCGTGGTGACCGTGACCCAGGACTCCTCCCTGCAGGACGGCGAGTTCATCTACAAG GTGAAGCTGCGCGGCACCAACTTCCCCTCCGACGGCCCCGTAATGCAGAAGAAGA CCATGGGCTGGGAGGCCTCCTCCGAGCGGATGTACCCCGAGGACGGCGCCCTGAA GGGCGAGATCAAGCAGAGGCTGAAGCTGAAGGACGGCGGCCACTACGACGCTGAG GTCAAGACCACCTACAAGGCCAAGAAGCCCGTGCAGCTGCCCGGCGCCTACAACG TCAACATCAAGTTGGACATCACCTCCCACAACGAGGACTACACCATCGTGGAACAGT ACGAACGCGCCGAGGGCCGCCACTCCACCGGCGGCATGGACGAGCTGTACAAGGA ATTTATGCAGACCATCAAATGCGTAGTCGTGGGTGATGGAGCAGTTGGCAAAACGTG TCTGCTCATCTCCTATACCACCAACAAATTTCCCAGCGAATACGTGCCTACGGTATTC GACAATTACGCGGTTACCGTCATGATTGGTGGTGAACCCTATACCTTGGGCCTGTTC GATACTGCAGGCCAAGAGGACTATGATCGCTTACGCCCTCTGTCGTATCCGCAAACC GACGTCTTTCTTGTGTGCTTTAGCGTTGTGTCTCCGAGTTCGTTCGAAAACGTGAAA GAGAAATGGGTACCGGAAATTACGCACCATTGTCCGAAAACTCCGTTTCTGCTGGTT GGCACACAGATCGATCTGCGCGATGATCCAAGCACCATTGAGAAACTTGCCAAGAAC AAACAGAAACCGATTACGCCAGAAACTGCGGAGAAATTAGCCCGTGATCTGAAAGCC GTCAAGTACGTGGAATGCTCAGCTTTGACACAGAAAGGGCTGAAGAATGTGTTTGAC GAAGCGATTCTGGCTGCGTTAGAACCGCCAGAACCGAAGAAAAGTCGTCGGTGTGT TCTCCTGTAA

### MSP1E3D1

Protein sequence MGSSHHHHHHENLYFQGSTFSKLREQLGPVTQEFWDNLEKETEGLRQEMSKDLEEVK AKVQPYLDDFQKKWQEEMELYRQKVEPLRAELQEGARQKLHELQEKLSPLGEEMRDR ARAHVDALRTHLAPYLDDFQKKWQEEMELYRQKVEPLRAELQEGARQKLHELQEKLSP LGEEMRDRARAHVDALRTHLAPYSDELRQRLAARLEALKENGGARLAEYHAKATEHLS TLSEKAKPALEDLRQGLLPVLESFKVSFLSALEEYTKKLNTQ*

#### DNA sequence

ATGGGCAGCAGCCATCATCATCATCATCATGAAAACCTGTATTTTCAGGGCAGCACCT TTAGCAAACTGCGTGAACAGCTGGGCCCGGTGACCCAGGAATTTTGGGATAACCTG GAAAAAGAAACCGAAGGCCTGCGTCAGGAAATGAGCAAAGATCTGGAAGAGGTGAA AGCGAAAGTGCAGCCGTATCTGGATGACTTTCAGAAAAAATGGCAGGAAGAGATGG AACTGTATCGTCAGAAAGTGGAACCGCTGCGTGCGGAACTGCAGGAAGGCGCGCG TCAGAAACTGCATGAACTGCAGGAAAAACTGAGCCCGCTGGGCGAAGAGATGCGTG ATCGTGCGCGTGCGCATGTGGATGCGCTGCGTACCCATCTGGCGCCGTATCTGGAT GACTTTCAGAAAAAATGGCAGGAAGAGATGGAACTGTATCGTCAGAAAGTGGAACCG CTGCGTGCGGAACTGCAGGAAGGCGCGCGTCAGAAACTGCATGAACTGCAGGAAA AACTGAGCCCGCTGGGCGAAGAGATGCGTGATCGTGCGCGTGCGCATGTGGATGC GCTGCGTACCCATCTGGCGCCGTATAGCGATGAACTGCGTCAGCGTCTGGCGGCCC GTCTGGAAGCGCTGAAAGAAAACGGCGGTGCGCGTCTGGCGGAATATCATGCGAAA GCGACCGAACATCTGAGCACCCTGAGCGAAAAAGCGAAACCGGCGCTGGAAGATCT GCGTCAGGGCCTGCTGCCGGTGCTGGAAAGCTTTAAAGTGAGCTTTCTGAGCGCG CTGGAAGAGTATACCAAAAAACTGAACACCCAGTAA

### Cloning methods

*E. coli* OneShot TOP10 (Invitrogen, Thermo Fisher Scientific, Waltham, USA) cells were used for propagation of all plasmids. All gene sequences created in this study were synthesized by Eurofins Genomics Europe in cloning vectors. Each gene was then amplified and subcloned into the expression vector using SLIC methods^3^. For fusion constructs with the C-terminal sequence of CAAX proteins, a long rigid linker^4^ (protein sequences in italics) was used, such as GST-KrasB, GST-Cdc42 and mCherry-Cdc42_CAAX_. The 3C protease recognition sequences were highlighted with underline.

### Preparation of T7 polymerase

Preparation of T7 polymerase was performed as described previously^5^. *E.coli* strain BL21(DE3) Star was transformed with plasmid pAR1219^6^, carrying the T7 ploymerase gene. 1 L of LB medium with antibiotics was inoculated with an overnight culture at 1:100 ratio. Cells were grown on a shaker at 37 ^°^C until Abs_600_ reached 0.6-0.8. T7 polymerase production was induced by addition of IPTG (1 mM final concentration in media). Cells were cultured for 5 hours and harvested by centrifugation at 8,000xg for 15 min at 4 ^°^C. Cell pellets were resuspended in 30 mL T7 buffer A (30 mM Tris-HCl, pH 8.0, 50 mM NaCl, 1 mM EDTA, 10 mM *β*-mercaptoethanol, 5% glycerol) and disrupted via single pass-through a French Press at 15,000 psi. Then, the cell lysate was clarified via centrifugation at 20,000xg for 30 min at 4°C and the supernatant was adjusted to a final concentration of 4% (w/v %) streptomycin sulfate. The sample was then centrifuged at 20,000xg for 30 min at 4 ^°^C. The resulting supernatant was filtered and loaded onto a 40 mL Q-sepharoase column pre-equilibrated with T7 buffer B (30 mM Tris-HCl, pH 8.0, 50 mM NaCl, 1 mM EDTA, 10 mM *β*-mercaptoethanol, 5% (v/v %) glycerol) and washed extensively with T7 buffer B after loading. T7 polymerase was then eluted using a linear gradient of 50-500 mM NaCl and 10 column volumes of T7 buffer B. Collected fractions were analyzed by SDS-PAGE. Fractions containing T7 polymerase (a predominant band around 90 kDa) were pooled and subsequently dialyzed against T7 buffer C (10 mM K_2_HPO_4_/KH_2_PO_4_, pH 8.0, 10 mM NaCl, 0.5 mM EDTA, 1 mM DTT, 5% (v/v %) glycerol) over night. Glycerol was added to a final concentration of 10% (v/v %) and the protein was concentrated to 3-4 mg mL^-1^ by ultrafiltration. Additional glycerol was added to a final concentration of 50% (v/v %), single-use aliquots were flash frozen using liquid nitrogen and stored at -80 ^°^C.

### Nanodisc preparation

The preparation of nanodiscs was performed according to previous protocols^2^. In brief, the MSP1E3D1 protein was purified via Ni-NTA column. After elution, the protein containing fractions were pooled with 10% glycerol (v/v) to prevent precipitation and were dialyzed over night against buffer (40 mM Tris-HCl, pH 8.0, 300 mM NaCl, 10% (v/v %) glycerol), including a buffer exchange after 2 hours. The resulting protein was centrifuged at 20,000xg for 15 min to remove precipitated proteins. The samples were then aliquoted, frozen in liquid nitrogen and stored at -80 °C until further usage. The nanodiscs used in this study were assembled via mixing of the following reagents: 25 μM MSP1E3D1, 1.6 mM DOPC (50 mM stock dissolved in 300 mM sodium cholate) (Avanti Polar lipids, Inc.), 0.4 mM DOPS (50 mM stock dissolved in 300 mM sodium cholate), 0.1% (v/v%) n- dodecylphosphocholine (DPC) in buffer (40 mM Tris-HCl, pH 8.0, 300 mM NaCl). The mixture was incubated for 1 hour at room temperature. Nanodisc-assembly was achieved by dialysis (1:500 volume ratio, buffer: 10 mM Tris-HCl, pH 8.0, 100 mM NaCl) using a Slide-A-Lyzer^TM^ (Thermo Fisher Science) at RT for 12 hours. The sample was again dialyzed for 24 hours at 4 ^°^C and centrifuged for 20 min at 20,000xg. The supernatant was collected and concentrated using an Amicon filter unit (10 kDa, MWCO, Millipore). Final concentration of nanodiscs should be above 0.5 mM, which corresponds to 1 mM MSP1E3D1. Concentrated nanodiscs were aliquoted, flash frozen in liquid nitrogen and stored in -80 ^°^C.

### Preparation of SLB chambers

#### Cleaning of Coverslips

24 × 24 mm #1.5 coverslips (Menzel) were piranha cleaned by adding 7 drops of sulfuric acid and two drops of 50 % hydrogen peroxide to the center of each coverslip. The reaction was incubated on the coverslips for at least 45 min before thoroughly rinsing with ultrapure water.

### Assembly of Reconstitution Chamber

The reaction chamber was formed through the attachment of a bisected 0.5 mL reaction tube onto cleaned coverslips, using an optical glue (Norland Optical Adhesive 68, Norland Products) that was cured under a UV lamp (365 nm) for 10 min.

## Supplementary Code

~~~
**Z-stack analysis macro code**
setPasteMode(“Copy”);
originalImage = getTitle();
getDimensions(width, height, channels, slices, frames);
newImage(originalImage + “_membraneSliceTimeSeries”, bitDepth+“-bit”, width, height, channels, 1, frames);
membraneSliceTimeSeries = getTitle();
for(t=1; t<=frames; t++) {
     selectWindow(originalImage);
     //Select the first channel, slice and frame t.
     Stack.setPosition(1,1,t);
     brightestMean = 0;
     brightestSlice = 1;
     for(i=1; i<slices; i++) {
        Stack.setPosition(1,i,t);
        getStatistics(area, mean);
        if(brightestMean<mean) {
            brightestMean = mean;
            brightestSlice = i;
        }
     }
     //print(brightestSlice); //test
     Stack.setPosition(1,brightestSlice,t);
     run(“Select All”);
     run(“Copy”);
     selectWindow(membraneSliceTimeSeries);
     Stack.setPosition(1,1,t);
     run(“Paste”);
     //print the second channel selectWindow(originalImage);
     Stack.setPosition(2,brightestSlice,t);
     run(“Select All”);
     run(“Copy”);
     selectWindow(membraneSliceTimeSeries);
     Stack.setPosition(2,1,t);
    run(“Paste”);
}
~~~

## References

1. Ganzinger, K. A., Schwille, P. More from less – bottom-up reconstitution of cell biology. J. Cell. Sci. 132, jcs227488 (2019).

2. Joyce, G. F., Szostak, J. W. Protocells and RNA Self-Replication. Cold Spring Harbor Perspect. Biol. 10, a034801 (2018).

3. Altenburg, W. J., Yewdall, N. A., Vervoort, D. F., van Stevendaal, M. H., Mason, A. F., van Hest, J. Programmed spatial organization of biomacromolecules into discrete, coacervate-based protocells. Nat. Commun. 11, 6282 (2020).

4. Dzieciol, A. J., Mann, S. Designs for life: protocell models in the laboratory. Chem. Soc. Rev. 41, 79–85 (2012).

5. Yoshida, A., Kohyama, S., Fujiwara, K., Nishikawa, S., Doi, N. Regulation of spatiotemporal patterning in artificial cells by a defined protein expression system. Chem. Sci. 10, 11064–11072 (2019).

6. Noireaux, V., Libchaber, A. A vesicle bioreactor as a step toward an artificial cell assembly. Proc. Natl. Acad. Sci. USA 101, 17669–17674 (2004).

7. Godino, E. et al. De novo synthesized Min proteins drive oscillatory liposome deformation and regulate FtsA-FtsZ cytoskeletal patterns. Nat. Commun. 10, 4969 (2019).

8. Exterkate, M., Caforio, A., Stuart, M. C., Driessen, A. J. Growing Membranes In Vitro by Continuous Phospholipid Biosynthesis from Free Fatty Acids. ACS Synth. Biol. 7, 153–165 (2018).

9. Schwille, P., et al. MaxSynBio - Avenues towards creating cells from the bottom up. Angew. Chem. Int. Ed. 57, 13382–13392 (2018).

10. Guindani, C., Caire da Silva, L., Cao, S., Ivanov, T., Landfester, K. Synthetic Cells: From Simple Bio-Inspired Modules to Sophisticated Integrated Systems. Angew. Chem. Int. Ed. 10.1002/anie.202110855 (2021).

11. Jia, H. Y., Heymann, M., Bernhard, F., Schwille, P., Kai, L. Cell-free protein synthesis in micro compartments: building a minimal cell from biobricks. New Biotechnol. 39, 199–205 (2017).

12. Berhanu, S., Ueda, T., Kuruma, Y. Artificial photosynthetic cell producing energy for protein synthesis. Nat. Commun. 10, 1325 (2019).

13. Libicher, K., Hornberger, R., Heymann, M., Mutschler, H.. In vitro self-replication and multicistronic expression of large synthetic genomes. Nat. Commun. 11, 904 (2020).

14. Niederholtmeyer, H., Chaggan, C., Devaraj, N. K. Communication and quorum sensing in non-living mimics of eukaryotic cells. Nat. Commun. 9, 5027 (2018).

15. Blanken, D., Foschepoth, D., Serrao, A. C., Danelon, C. Genetically controlled membrane synthesis in liposomes. Nat. Commun. 11, 4317 (2020).

16. Jaroentomeechai, T., et al. Single-pot glycoprotein biosynthesis using a cell-free transcription-translation system enriched with glycosylation machinery. Nat. Commun. 9, 2686 (2018).

17. van Nies, P., Westerlaken, I., Blanken, D., Salas, M., Mencia, M., Danelon, C. Self- replication of DNA by its encoded proteins in liposome-based synthetic cells. Nat. Commun. 9, 1583 (2018).

18. Godino, E., et al. Cell-free biogenesis of bacterial division proto-rings that can constrict liposomes. *Commun*. Biol. 3, 539 (2020).

19. Rojas, A. M., Fuentes, G., Rausell, A., Valencia, A. The Ras protein superfamily: Evolutionary tree and role of conserved amino acids. J. Cell Biol. 196, 189–201 (2012).

20. Zinatizadeh, M. R., et al. The Role and Function of Ras-association domain family in Cancer: A Review. Genes Dis. 6, 378–384 (2019).

21. Cerione, R. A. Cdc42: new roads to travel. Trends Cell Biol. 14, 127–132 (2004).

22. Woods, B., Lew, D. J. Polarity establishment by Cdc42: Key roles for positive feedback and differential mobility. Small GTPases 10, 130–137 (2019).

23. Zhang, F. L., Casey, P. J. Protein prenylation: Molecular mechanisms and functional consequences. Annu. Rev. Biochem. 65, 241–269 (1996).

24. Wang, M., Casey, P. J. Protein prenylation: unique fats make their mark on biology. Nat. Rev. Mol. Cell Biol. 17, 110–122 (2016).

25. Casey, P. J., Seabra, M. C. Protein prenyltransferases. J. Biol. Chem. 271, 5289–5292 (1996).

26. Long, S. B., Casey, P. J., Beese, L. S. Reaction path of protein farnesyltransferase at atomic resolution. Nature 419, 645–650 (2002).

27. Taylor, J. S., Reid, T. S., Terry, K. L., Casey, P. J., Beese, L. S. Structure of mammalian protein geranylgeranyltransferase type-I. EMBO J. 22, 5963–5974 (2003).

28. Shirakawa, R. et al. A SNARE geranylgeranyltransferase essential for the organization of the Golgi apparatus. EMBO J. 39, e104120 (2020).

29. Moores, S. L. et al. Sequence Dependence of Protein Isoprenylation. J. Biol. Chem. 266, 14603–14610 (1991).

30. Armstrong, S. A., Seabra, M. C., Sudhof, T. C., Goldstein, J. L., Brown, M. S. Cdna Cloning and Expression of the Alpha-Subunit and Beta-Subunit of Rat Rab Geranylgeranyl Transferase. J. Biol. Chem. 268, 12221–12229 (1993).

31. Rak, A., Pylypenko, O., Niculae, A., Pyatkov, K., Goody, R. S., Alexandrov, K. Structure of the Rab7 : REP-1 complex: Insights into the mechanism of rab prenylation and choroideremia disease. Cell 117, 749–760 (2004).

32. Suzuki, T., et al. Protein prenylation in an insect cell-free protein synthesis system and identification of products by mass spectrometry. Proteomics 7, 1942–1950 (2007).

33. Nguyen, U. T. et al. Exploiting the substrate tolerance of farnesyltransferase for site-selective protein derivatization. ChemBioChem 8, 408–423 (2007).

34. Dursina, B., et al. Identification and specificity profiling of protein prenyltransferase inhibitors using new fluorescent phosphoisoprenoids. J. Am. Chem. Soc. 128, 2822–2835 (2006).

35. Hannoush, A. N., Sun, J. The chemical toolbox for monitoring protein fatty acylation and prenylation. Nat. Chem. Biol. 6, 498–506 (2010).

36. Kho, Y. et al. A tagging-via-substrate technology for detection and proteomics of farnesylated proteins. Proc. Natl. Acad. Sci. USA 101, 12479–12484 (2004).

37. Berry, A. F. et al. Rapid Multilabel Detection of Geranylgeranylated Proteins by Using Bioorthogonal Ligation Chemistry. ChemBioChem 11, 771–773 (2010).

38. Ramm, B., Heermann, T., Schwille, P. The E. coli MinCDE system in the regulation of protein patterns and gradients. Cell Mol. Life Sci. 76, 4245–4273 (2019).

39. Harrington, L., Fletcher, J. M., Heermann, T., Woolfson, D. N., Schwille, P. De novo design of a reversible phosphorylation-dependent switch for membrane targeting. Nat. Commun. 12, 1472 (2021).

40. Zhang, F. L., et al. Cdna Cloning and Expression of Rat and Human Protein Geranylgeranyltransferase Type-I. J. Biol. Chem. 269, 3175–3180 (1994).

41. Boland, C. et al. Cell-free expression and in meso crystallisation of an integral membrane kinase for structure determination. Cell Mol. Life Sci. 71, 4895–4910 (2014).

42. Foshag, D. et al. The E. coli S30 lysate proteome: A prototype for cell-free protein production. New Biotechnol. 40, 245–260 (2018).

43. Henrich, E., Hein, C., Dötsch, V., Bernhard, F. Membrane protein production in Escherichia coli cell-free lysates. FEBS Lett. 589, 1713–1722 (2015).

44. Denisov, I. G., Sligar, S. G. Nanodiscs for structural and functional studies of membrane proteins. Nat. Struct. Mol. Biol. 23, 481–486 (2016).

45. Williams, C. L. The polybasic region of Ras and Rho family small GTPases: a regulator of protein interactions and membrane association and a site of nuclear localization signal sequences. Cell. Signal. 15, 1071–1080 (2003).

46. Tnimov, Z. et al. Quantitative analysis of prenylated RhoA interaction with its chaperone, RhoGDI. J. Biol. Chem. 287, 26549–26562 (2012).

47. Hoffman, G. R., Nassar, N., Cerione, R. A. Structure of the Rho family GTP-binding protein Cdc42 in complex with the multifunctional regulator RhoGDI. Cell 100, 345–356 (2000).

48. Snyder, J. T. et al. Structural basis for the selective activation of Rho GTPases by Dbl exchange factors. Nat. Struct. Biol. 9, 468–475 (2002).

49. Quevedo, C. E. et al. Small molecule inhibitors of RAS-effector protein interactions derived using an intracellular antibody fragment. Nat. Commun. 9, 3169 (2018).

50. Kristelly, R., Gao, G., Tesmer, J. J. Structural determinants of RhoA binding and nucleotide exchange in leukemia-associated Rho guanine-nucleotide exchange factor. J. Biol. Chem. 279, 47352–47362 (2004).

51. Cruz-Migoni, A. et al. Structure-based development of new RAS-effector inhibitors from a combination of active and inactive RAS-binding compounds. Proc. Natl. Acad. Sci. USA 116, 2545–2550 (2019).

52. Bendezu, F. O., Vincenzetti, V., Vavylonis, D., Wyss, R., Vogel, H., Martin, S. G. Spontaneous Cdc42 Polarization Independent of GDI-Mediated Extraction and Actin-Based Trafficking. Plos. Biol. 13, e1002097 (2015).

53. Lamas, I., Merlini, L., Vjestica, A., Vincenzetti, V., Martin, S. G. Optogenetics reveals Cdc42 local activation by scaffold-mediated positive feedback and Ras GTPase. Plos. Biol. 18, e3000600 (2020).

54. Gillette, W. K. et al. Farnesylated and methylated KRAS4b: high yield production of protein suitable for biophysical studies of prenylated protein-lipid interactions. Sci. Rep. 5, 1–13 (2015).

55. Manolaridis, I. et al. Mechanism of farnesylated CAAX protein processing by the intramembrane protease Rce1. Nature 504, 301–305 (2013).

56. Mazhab-Jafari, M. T. et al. Oncogenic and RASopathy-associated K-RAS mutations relieve membrane-dependent occlusion of the effector-binding site. Proc. Natl. Acad. Sci. USA 112, 6625–6630 (2015).

57. Mazhab-Jafari, M. T. et al. Membrane-Dependent Modulation of the mTOR Activator Rheb: NMR Observations of a GTPase Tethered to a Lipid-Bilayer Nanodisc. J. Am. Chem. Soc. 135, 3367–3370 (2013).

58. Hobbs, G. A., Der, C. J., Rossman, K. L. RAS isoforms and mutations in cancer at a glance. J. Cell Sci. 129, 1287–1292 (2016).

59. Wheeler, A. P., Ridley, A. J. Why three Rho proteins? RhoA, RhoB, RhoC, and cell motility. Exp. Cell Res. 301, 43–49 (2004).

60. Zawada, J. F. et al. Microscale to Manufacturing Scale-up of Cell-Free Cytokine Production-A New Approach for Shortening Protein Production Development Timelines. Biotechnol. Bioeng. 108, 1570–1578 (2011).

61. Ramm, B. et al. The MinDE system is a generic spatial cue for membrane protein distribution in vitro. Nat. Commun. 9, 3942 (2018).

62. Jia, H. Y., Kai, L., Heymann, M., Garcia-Soriano, D. A., Hartel, T., Schwille, P. Light- Induced Printing of Protein Structures on Membranes in Vitro. Nano Lett. 18, 7133–7140 (2018).

63. Etienne-Manneville, S. Cdc42 - the centre of polarity. J. Cell Sci. 117, 1291–1300 (2004).

64. Johnson, D. I. Cdc42: An essential Rho-type GTPase controlling eukaryotic cell polarity. Microbiol. Mol. Biol. Rev. 63, 54–105 (1999).

65. Kalinin, A., et al. Expression of mammalian geranylgeranyltransferase type-II in Escherichia coli and its application for in vitro prenylation of Rab proteins. Protein Expr. Purif. 22, 84–91 (2001).

66. Kigawa, T., et al. Preparation of Escherichia coli cell extract for highly productive cell-free protein expression. J. Struct. Funct. Genomics 5, 63–68 (2004).

67. Kai, L., et al. Systems for the cell-free synthesis of proteins. Methods Mol. Biol. 800, 201–225 (2012).

68. Schindelin, J., et al. Fiji: an open-source platform for biological-image analysis. Nat. Methods 9, 676–682 (2012).

69. Glock, P. et al. Stationary Patterns in a Two-Protein Reaction-Diffusion System. ACS Synth. Biol. 8, 148–157 (2019).

70. Ramm, B., Glock, P., Schwille, P. In Vitro Reconstitution of Self-Organizing Protein Patterns on Supported Lipid Bilayers. JoVE e58139, https://doi.org/10.3791/58139 (2018).

## Supplementary References

1. Dursina, B. et al. Identification and specificity profiling of protein prenyltransferase inhibitors using new fluorescent phosphoisoprenoids. J. Am. Chem. Soc. 128, 2822–2835 (2006).

2. Roos, C. et al. Co-translational association of cell-free expressed membrane proteins with supplied lipid bilayers. Mol. Membr. Biol. 30, 75–89 (2013).

3. Scholz, J., Besir, H., Strasser, C., Suppmann, S. A new method to customize protein expression vectors for fast, efficient and background free parallel cloning. BMC Biotechnol. 13, 12 (2013).

4. Chen, X. Y., Zaro, J. L., Shen, W. C. Fusion protein linkers: Property, design and functionality. Adv. Drug. Deliver. Rev. 65, 1357–1369 (2013).

5. Kai, L. et al. Systems for the cell-free synthesis of proteins. Methods Mol. Biol. 800, 201–225 (2012).

6. Li, Y., Wang, E. D., Wang, Y. L. A modified procedure for fast purification of T7 RNA polymerase. Protein Expres. Purif. 16, 355–358 (1999).

